# Functional cell type diversity in the *Drosophila* lamina is established by a single transcription factor dFezf, controlling a large repertoire of ion channels and receptors

**DOI:** 10.64898/2026.11.19.766796

**Authors:** Katja Sporar, Neel Wagh, Junaid Akhtar, Ivan J Santiago, Marko Ilić, Matthew Pecot, Marion Silies

## Abstract

Neuronal cell types in the brain can show remarkable diversity in their functional properties. In the visual system, neurons at different levels of hierarchy show diverse kinetics, leading to the encoding of different features of the visual scene. How this behaviorally relevant diversification is achieved is not understood. Here, we address this question in a simple system, studying two visual cell types in *Drosophila* that diversify their responses despite receiving the same photoreceptor input to encode two distinct basic visual features, contrast and luminance. We first show that this cell type diversity is mostly established by cell-intrinsic mechanism. While cell-type specific transcriptome analysis delivers promising receptor and channel candidates, genetic manipulation of these differentially regulated genes fails to identify individual channels or receptors that are necessary or sufficient for the functional differences between luminance-sensitive L3 neurons and contrast-sensitive L2 neurons. However, loss of the L3-specific transcription factor dFezf makes L3 neurons transient and contrast-sensitive. Interestingly, dFezf controls the cell-type-specific expression of several ion channels and receptors that we initially found to be differentially expressed between L2 and L3. Together, this argues for a degenerate system in which multiple genes jointly shape cell type-specific functional properties. Our finding that a single transcription factor regulates the expression of such a set of genes that can be relevant for function suggest that similar regulation might also be in place in other degenerate systems, which exist widely across cell types, brain regions, and species.

## Introduction

In the nervous system, neuronal cell types are defined by specific combinations of morphological, anatomical, and physiological features. These features can be very diverse between cell types, even if the cell types are at the same layer of processing. This diversity is important for their function within neural circuits. In the visual system, for example, functionally diverse cell types are essential to compute motion, or to extract fundamentally different features from the visual scene, such as luminance and contrast. Synapse-level connectomes provide information about which cell type is communicating with whom, and via how many synapses. However, in some cases, functionally relevant cell-type diversity is established even when distinct cell types share a common input. Here, differences in gene expression can establish diversity. While it is well understood how distinct cell types emerge through the serial expression of different transcription factors, the factors that further diversify cell types in terms of their functional properties are less well studied. Here, we address this question using the example of functional diversification of cell types postsynaptic to photoreceptors in the *Drosophila* visual system.

In the visual system, temporal diversity exists in neuronal cell types at different levels of hierarchy. In both vertebrates and invertebrates, the neurons directly postsynaptic to photoreceptors - bipolar cells in vertebrates or lamina neurons in invertebrates - already show a repertoire of temporal dynamics. These distinct properties enable them to encode fundamentally different features of the visual scene, namely contrast and luminance. This temporal cell-type diversity then increases postsynaptically, as illustrated by the wide repertoire of temporal filtering properties in the third order visual neurons: the retinal ganglion cells in the vertebrate retina, or medulla neurons in the fly. These specific functional properties allow classifying neurons into cell types in the first place ^1–4^. The distinct temporal kinetics are important for function in multiple ways. For example, differential temporal filtering of the visual input is a prerequisite for the computation of motion direction ^5–7^. Furthermore, temporal cell type diversity in the *Drosophila* visual system has been shown to reflect natural stimulus statistics, such that temporally distinct filters encode naturalistic inputs more efficiently than circuits composed of more similar filters ^8^. Thus, the emergence of temporal diversity plays a critical functional role along the hierarchy of visual pathways. What differentiates functionally relevant properties within such cell types, including their temporal features, remains unknown in most systems.

Temporal cell-type diversity originates at the first synapse between the photoreceptors and their postsynaptic partners. These second-order visual neurons range from transient neurons with biphasic temporal filters that respond to changes in their visual input and thus encode contrast, to sustained neurons with monophasic temporal filters that report the luminance of the visual input ^9–12^. In the vertebrate retina, bipolar cells downstream of photoreceptors show a diversity of temporal properties ^2,3,9,13^. While most bipolar cells compute contrast, the sustained component of the rod bipolar cell synapse has been shown to encode luminance ^9^. In *Drosophila*, the lamina neurons L1, L2, and L3 are postsynaptic to photoreceptors R1-R6 at nanoscale proximity at so-called tetrad synapses ^14^. R1-R6 are broad-band sensitive photoreceptors that feed information into achromatic visual pathways. Of the lamina neurons, L2 neurons are characterized by a biphasic filter and are sensitive to contrast, whereas L3 neurons show a monophasic filter and are sensitive to luminance ^11,15,16^. L1 has a biphasic filter and closely resembles L2, but with an additional luminance-sensitive component ^15,16^. Both luminance and contrast information are functionally relevant in downstream circuitry. The presence of luminance information postsynaptic to photoreceptors has been shown to support vision in dynamically changing light conditions, typically encountered when navigating in natural environments. Under such conditions, the L3 luminance signal provides the basis for rapid luminance gain control. L3 is thus required for robust contrast computation when background illumination is rapidly changing ^10,16–18^. How this functionally relevant temporal diversity arises is not understood in any system.

In the vertebrate retina, photoreceptors release glutamate, and differential expression of ionotropic glutamate receptors, particularly AMPA and kainate receptors might contribute to temporal diversity among OFF bipolar cells ^19–21^. In the *Drosophila* visual system, photoreceptors release histamine as the primary neurotransmitter. However, all lamina neurons that receive direct input from R1-R6, express the same histamine-gated chloride channel Ort (Outer rhabdomeres transientless), arguing that the receptor itself does not add to the functional cell type diversity. A second histamine-gated chloride channel, HisCl1, is thought to be absent from lamina neurons, but instead is expressed on glial cells and photoreceptors R7, R8, involved in color-vision ^22–25^. Voltage-gated potassium (K_v_) channels have been proposed as active regulators of membrane properties that underlie different properties and mediate distinct currents in lamina neurons of various insects ^26–28^. A pharmacological and genetic analysis of *Drosophila* K_v_ channels revealed the A-type K_v_ channels Shaker and Shal function to enhance responses and sharpen kinetics in L2, but the prominent cell-type specific differences between L2 and L3 could not be explained ^28^. Circuit interaction (chemical or electrical synaptic interactions) could also contribute, and a recent study showed that L2 filters are sharpened by recurrent synaptic feedback ^29^. However, despite several efforts to identify cell-type specific differences, the remarkable functional diversity of L1/L2 vs. L3 neurons remains unexplained.

The possibility emerges that no single molecular component can alone explain lamina neuron temporal diversity. In fact, many examples exist in neuroscience, where neuronal properties are jointly regulated by a wider range of components. In such degenerate systems, several solutions are implemented that all result in the same functional properties. In systems as diverse as neurons of the crab stomatogastric ganglion or mouse hippocampal neurons, different sets of ion channels result in the same neuronal firing patterns ^30–34^. These systems are robust to perturbations of individual components. At the same time, the set of ion channels contributing to the same function is not random, but gene expression patterns often covary or are correlated in models, arguing for joint regulation ^30,35^. Homeostatic regulation and activity-dependent mechanisms have been suggested to play a role in controlling the structured expression ^30,35^. Furthermore, one can envision that coordinated control of gene expression via a common transcription factor contributes to this covariation. However, if such overarching regulation exists and contributes to the emergence of temporal cell type diversity in the visual system is not known.

Here, we focus on temporally diverse neurons of the fly visual system, concentrating on two cell types whose vastly different temporal properties are diversified downstream of the same photoreceptor input. Cell-type-specific transcriptome analysis reveals L2- and L3-specific candidate genes that could mediate their distinct physiological properties. However, functional testing of candidate genes such as the second histamine-gated chloride channel, HisCl1, and many other channel or receptor candidates, resulted in weak or no phenotypes. We thus decided to investigate a molecule class that can potentially regulate many genes together, through joint transcriptional regulation. We found that a single L3-specific transcription factor, dFezf (Erm), is required to mediate the temporal dynamics of L3 and that cell-type-specific loss of dFezf in L3 results in transient responses to visual stimuli that encode contrast, rather than luminance. Furthermore, transcriptome analysis of *dFezf* mutant L3 cells showed that dFezf acts as a regulator of many channels and receptors that differ between L2 and L3. Together, our results argue that temporal cell type diversity in the lamina is established by a degenerate set of channels and receptors, which are jointly regulated by a cell-type specific transcription factor.

## Results

### Distinct kinetics of L2 and L3 responses emerge in their dendrites

Lamina neurons L2 and L3 respond to fundamentally different features of the visual scene, despite receiving joint R1-R6 photoreceptor input. When recording cell type-specific in vivo calcium responses, L2 neurons show transient increases in calcium signals in response to light decrements (OFF steps) that quickly return to one common baseline, and have previously been shown to encode contrast (**Fig. 1A-C**) ^10,15,16^. Calcium signals in L3 neurons are sustained and encode luminance, such that L3 responses are higher when the stimulus is darker, and lower when the stimulus is brighter (**Fig. 1A-C**) ^10,11,16^. These properties could emerge at the photoreceptor-to-lamina neuron synapse, emerge through cell-intrinsic properties of L2 and L3, or even be shaped by further circuit mechanisms. Lamina neuron calcium signals have typically been recorded from their axon terminals. To understand where these distinct lamina neuron response properties first arise, we recorded calcium signals along the L2 and L3 neurons: in their dendrites, along their axons, and in their axon terminals (**Fig. 1D-G**). Transient L2 responses already differed from sustained L3 responses in lamina neuron dendrites. While L3 responses stayed similarly sustained along the different neuronal compartments, L2 calcium signals became more transient along the neurite with the most transient responses recorded in their presynaptic axon terminals (**Fig. 1D-G**).

**Figure 1.**
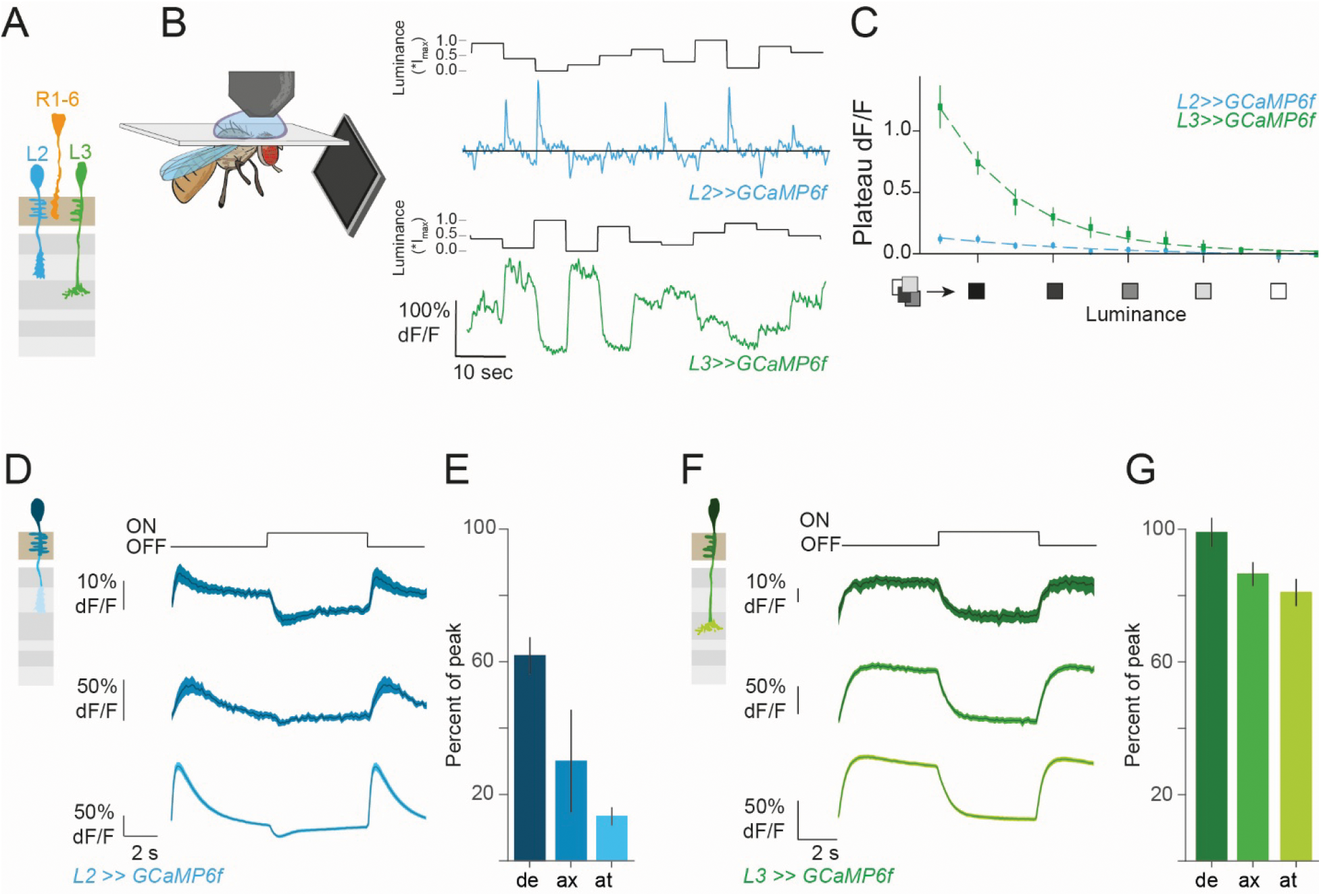
L2 transient vs L3 sustained response properties are established in lamina neuron dendrites. **(A)** Schematic of R1-R6 photoreceptors projecting to the lamina neurons L2 and L3 dendrites in the lamina. **(B)** Schematic of the in vivo 2 photon calcium imaging set-up. Calcium response of a single L2 or L3 axon terminal to a stimulus of randomly varying luminance, I_max_ was 1.<u>9</u>3∗10^5^ photons∗s^−1^∗receptor^−1^. **(C)** Quantification of the plateau responses (last 500 ms of the epoch) of all steps for a given luminance. **(D-G)** Average calcium (GCaMP6f) signals for L2 neurons (**D,E**) or L3 (**F,G**) from three different regions: dendrite, axon, and axon terminal, to 5 s full field flashes (**D,F**) and quantification of the sustenance as percentage of peak response retained during the plateau phase (**E,G**). Traces show mean ±SEM. Sample size, L2: n=9 (66), L3: n=6 (93) for (**C**), L2 dendrites: n=9 (43), L2 axon: n=7 (23), L2 axon terminals: n=23 (130) for (**D, E**), L3 dendrites: n=4 (18), L3 axon: n=14 (66), L3 axon terminals: n=22 (224) for (**F, G**). Sample sizes are given as number of flies (number of cells).

### R1-R6 but not R8 photoreceptors are sufficient inputs for L2 and L3 function

Lamina neurons L2 and L3 receive major anatomical input from R1-R6 in each lamina cartridge, but photoreceptors R8 functionally contributes to the motion detection pathway in flies (Wardill et al. 2012) and are less adapting ^36^, arguing that they could contribute to more sustained responses in some lamina neurons. To understand which photoreceptors predominantly shape the L2 and L3 responses (**Fig. 2A**), we analyzed the spectral efficiencies of L2 and L3 and compared them to photoreceptor properties. For visual stimulation, we used a multispectral synthesizer based on monochromatic LEDs and a diffraction grating ^37^ and presented light flashes of wavelengths spanning from UV to red to flies expressing GCaMP6f in either L2 or L3 neurons (**Fig. 2B**). The spectral efficiency curves of L2 and L3 axon terminals, obtained from the neurons’ OFF responses, showed similar sensitivities, both peaking at approximately 500nm (cyan). This closely matches the Rh1 spectral sensitivity ^38^ (**Fig. 2C**), arguing that both L2 and L3 predominantly rely on functional R1-R6 input. L3 responses showed a second peak at around 437nm, suggesting a potential contribution of R8 to L3 responses (**Fig. 2C**). To further test for the contribution of different photoreceptor inputs, we measured L2 and L3 calcium signals in flies in which either only R1-R6 or only R8 phototransduction was intact. To achieve this, we used *norpA* (*no receptor potential A*) mutants, which are blind, and then rescued *norpA* specifically in either R1-R6 or R8 (**Fig. 2D-I**). Control L2 axon terminals responded more transiently to the two OFF steps (gray-to-OFF, ON-to-gray) than L3 (**Fig. 2E,F**). Additionally, control L3 responded predominantly to the gray-to-OFF step, reflecting the nonlinear luminance dependence of L3 neurons (**Fig, 2E,F**). Both L2 and L3 responses in R1-R6 *norpA* rescues were indistinguishable from controls (**Fig. 2D-F**), showing that R1-R6 can drive normal L2 and L3 responses. Conversely, *norpA* expression in R8 did not restore normal L2 or L3 responses (**Fig. 2G-I**). Taken together, these results demonstrate that R1-R6 is the predominant functional input to both L2 and L3.

**Figure 2.**
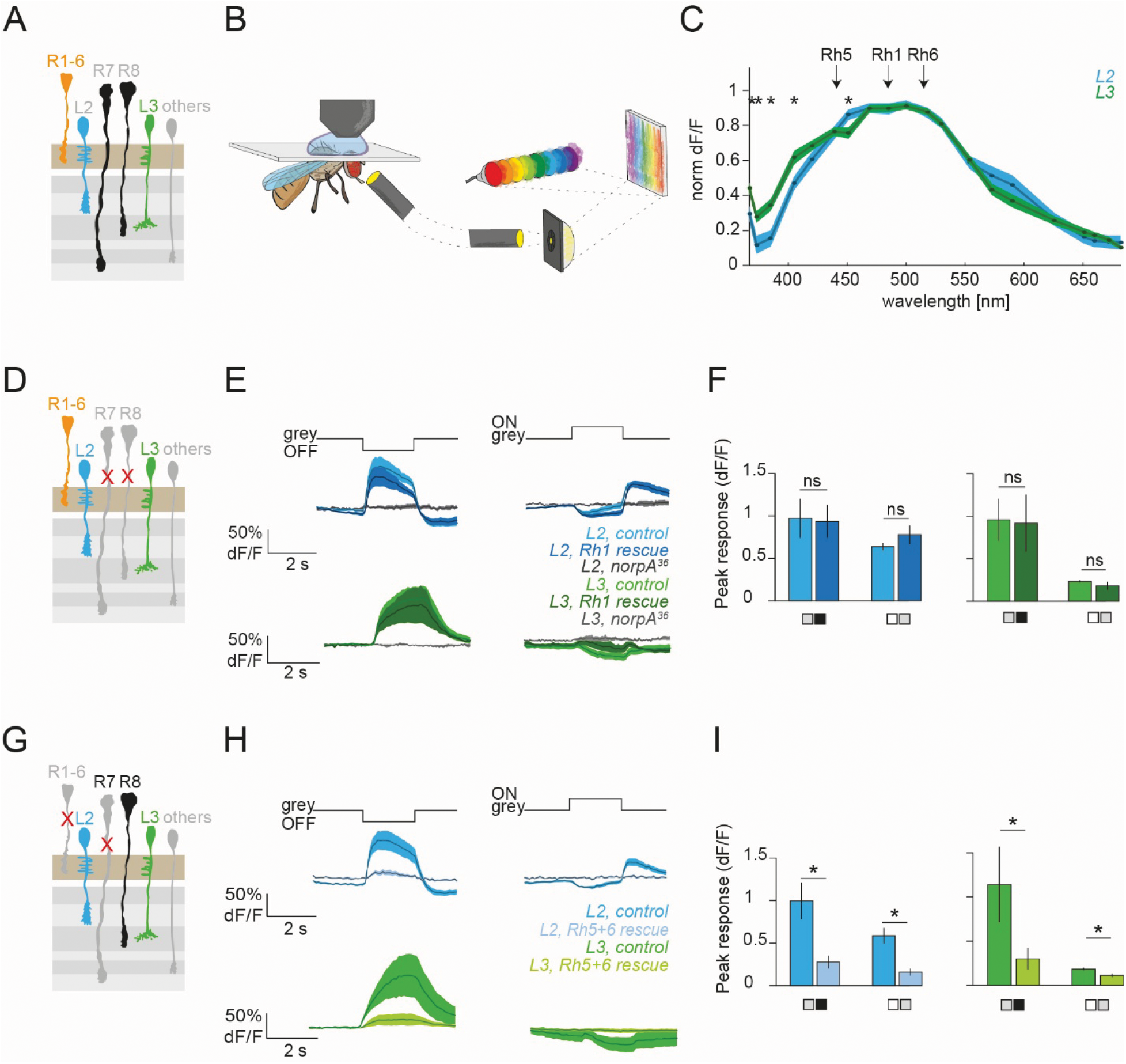
R1-R6 but not R8 photoreceptors are sufficient for normal L2 and L3 responses. **(A)** Schematic of achromatic R1-R6 and color R7 and R8 photoreceptors projecting to the lamina. R1-R6 project to L2 and L3 dendrites in lamina at the tetrad synapse, whereas R7 and R8 project to medulla. **(B)** Schematic of the LedSynth optical setup. Light from 20 monochromatic LEDs is directed onto the fly’s eye via a planar reflective diffraction grating, a launching lens, and a light guide. **(C)** Spectral efficiency curves for L2 and L3 neurons. Traces for peak (mean ± SEM) responses to the light offset of different wavelengths for L2 and L3. The peak sensitivities of the R1-R6 rhodopsin Rh1 (478 nm) and the R8 rhodopsin Rh5 (437 nm) and Rh6 (508 nm) are indicated by arrows. **(D)** Schematic for *norpA^36^* mutant rescue reestablishing only R1-R6 to L2-L3 connection. **(E)** Average calcium (GCaMP6f) signals from axon terminals for L2 and L3 in control, Rh1 rescue and *norpA^36^* mutant to the full-field flashes from an intermediate grey (ON-to-grey and grey-to-OFF). **(F)** Bar plots showing the quantification of the peak responses of control and R1-R6 rescued animals to the two OFF steps shown in (**E**): ON-to-grey and grey-to-OFF. (**G-I**) Same as (**D-F**) for R5+6 rescue, not showing the *norpA^36^* mutant again. *p<0.05, tested with two-tailed Student t tests (**C**), ns, not significant, *p<0.05, tested with two-tailed Student t tests (**I**). Sample size, L2: n= 4 (17) and L3: n= 8 (28) for (**C**), *L2, control*: n =5 (45), *L2, Rh1 rescue*: n=10 (73), *L2, norpA^36^: n =5(37), L3, control*: n=6 (50), *L3, Rh1 rescue*: n=5 (43), *L3, norpA^36^:* n=6 (48) for (**E,F**), *L2, control*: n=7 (56), *L2, R5+6 rescue*: n= 9 (50), *L3, control*: n=4 (28), *L3, R5+6 rescue*: n=16 (129 cells) for (**H, I**). Sample sizes are given as number of flies (number of cells).

To understand what transformations are occurring at the R1-R6 to lamina neuron synapses, we next recorded luminance and contrast responses of R1-R6 photoreceptors under the same stimulus conditions used to characterize L2 and L3 responses. We used a *ninaE-GCaMP6f* promotor fusion to measure the calcium response in the R1-R6 axon terminals. To test for luminance vs contrast sensitivity, we employed a well-established luminance-contrast discrimination stimulus ^9,10^. Here, two consecutive 3s OFF steps that followed an adapting bright period varied in both luminance and contrast (A step) or were fixed at 25% contrast but varied in luminance (B step) (**Fig. S1A**). Hence, a contrast-sensitive neuron will respond similarly to the different 25% contrast B steps (regardless of luminance), whereas a luminance-sensitive neuron will respond similarly to nearby luminance of the A and B steps (regardless of contrast). R1-R6 shows sustained calcium responses to all stimuli and responded similarly to close-by luminance of the A and B steps, whereas the responses to the six 25% contrast B steps differed (**Fig. S1A-C**). This revealed that R1-R6 calcium signals are sensitive to luminance in the given stimulus regime. Both here, and when using a stimulus of 10 s duration with randomly varying luminance, R1-R6 responses scaled with luminance (**Fig. S1D,E**). Taken together, calcium signals in R1-R6 axon terminals carry luminance information and - within the tested luminance range - encode luminance more linearly than L3, which shows enhanced sensitivity towards dark luminance. With this, both L2 and L3 differentially transform common photoreceptor input to obtain their distinct contrast and luminance sensitivities and hence extract different features of the visual environment.

### Lamina neuron properties are mostly controlled by cell-autonomous mechanisms, and only little by circuit interactions

To understand whether L2 and L3 obtain their distinct response profiles through circuit mechanisms downstream of R1-R6, or through cell-autonomous mechanisms, we genetically isolated L2 or L3 neurons from most circuit interactions (**Fig. 3A,B**). In *Drosophila*, histamine is the main neurotransmitter used by the photoreceptors, with the histamine-gated channel Ort being the major neuronal receptor ^23–25^. We used *ort* mutants, which have strong deficits in visual responses ^24,39^, and selectively restored *ort* expression in either L2 or L3, such that only photoreceptor-to-L2 synapses are intact in an L2-ort-rescue and only photoreceptor-to-L3 synapses are intact in an L3-ort-rescue. In *ort* mutants, L2 and L3 neurons exhibit a marked reduction in, but non-zero, responses to full-field flashes. Restoring ort expression in either cell types restored transient (L2, **Fig. 3A**) or sustained (L3, **Fig. 3B**) responses. Given the slight changes in response kinetics, we further investigated luminance encoding using full-field light flashes of randomly varying luminance. Whereas control L2 neurons, which responded to the OFF steps transiently and returned to a fixed baseline, L2 neurons in the *L2-ort-rescue* showed the emergence of a sustained plateau response along with the transient peak response (**Fig. 3C**). The plateau response in these neurons encoded luminance non-linearly, with the highest response at the darkest luminance, similar to L3 neurons (**Fig. 3C,E**). When we instead rescued *ort* expression in L3 neurons, L3 neurons maintained the same sustained response profiles as in controls (**Fig. 3 D,E**). Taken together, our data argues that L3 neuronal responses are not shaped by circuit interactions. Whereas lateral circuitry appears to be able to eliminate a luminance-sensitive baseline from L2 neurons, the transient initial response of L2 neurons is not affected by circuit properties. Together, these findings argue that most of the L2 vs L3 differences are mediated by cell-autonomous mechanisms.

**Figure 3.**
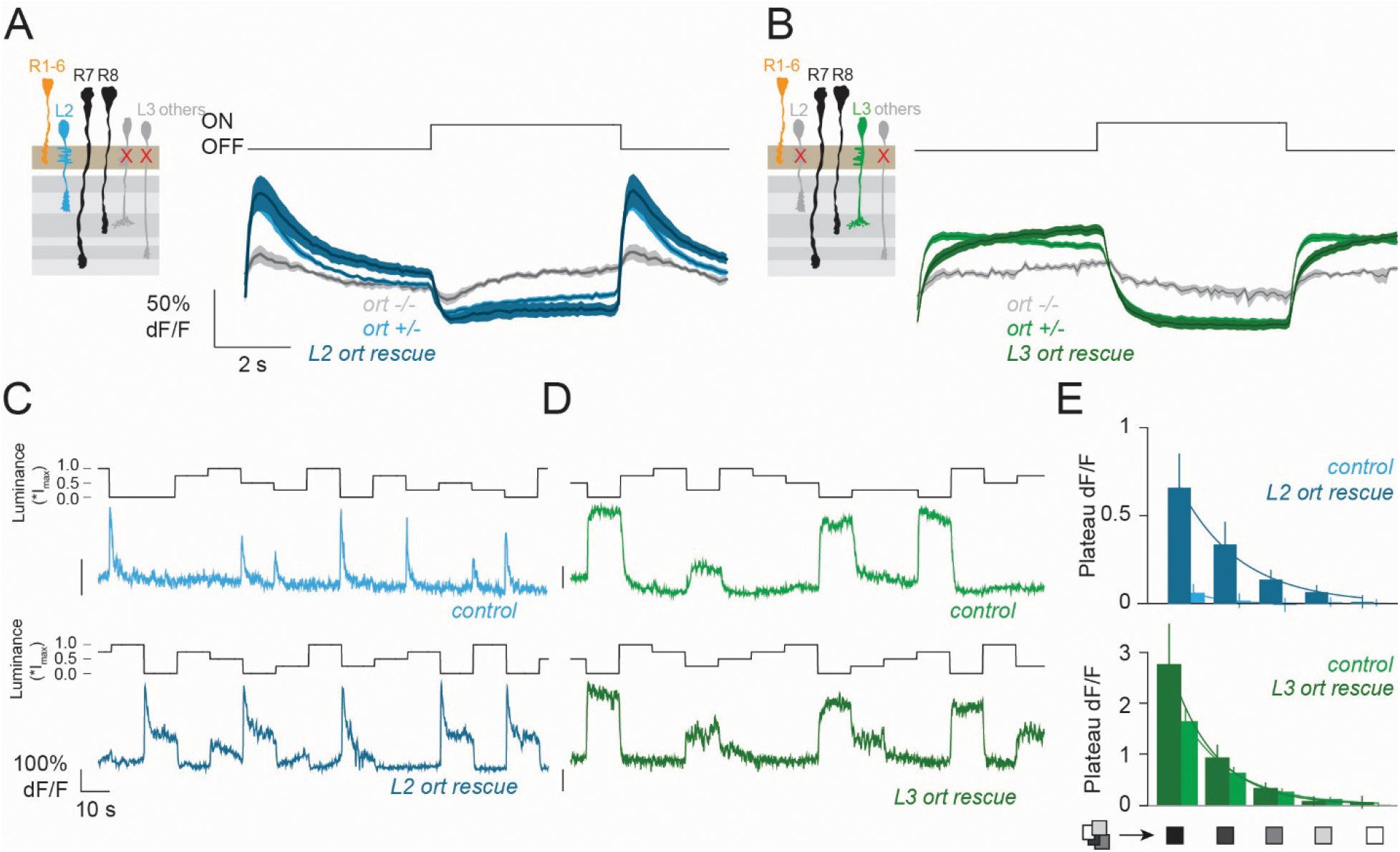
In the absence of circuit interactions, L2 neurons retain a luminance-sensitive baseline. **(A,B)** L2 (A) or L3 (B) *ort* rescues. Left; Schematics of the experiment. Right: Average calcium (GCaMP6f) signals from L2 (**A**) or L3 (**B**) axon terminals for *ort* heterozygous control, *ort* mutant and *ort* rescue to 5 s full field flashes. **(C)** Calcium response of a single L2 control and L2 *ort* rescue axon terminal to a stimulus of randomly varying luminance. The maximum screen luminance Imax was 1.87∗10^5^ photons∗s^−1^∗receptor^−1^. **(D)** Same as (**C**) for L3 control and L3 *ort* rescue. **(E)** Plateau responses (last 2 s of the epoch) of all steps for a given luminance. Sample size, *L2 ort+/-*: n=14 (83), *L2 ort-/-*: n=6 (26), *L2 ort rescue*: n=9 (74) for (**A**), *L3 ort+/-*: n=18 (193), *L3 ort-/-*: n=6 (39), *L3 ort* rescue: n=11 (145) for (**B**), *L2 control*: n=16 (140), *L2 ort rescue*: n=13 (143), *L3 control*: n=10 (73), *L3 ort rescue*: n=12 (91) for (**E**). Sample sizes are given as number of flies (number of cells).

### HisCl1 is as a second histamine receptor in lamina neurons, but is not responsible for cell-type specific response properties

To identify candidate genes that cell-autonomously establish the functional response properties of L2 and L3, we set out to explore genes that are differentially expressed in these cell types. Starting with existing cell-type specific transcriptome data ^40^, we found that this dataset contained genes not plausibly expressed in lamina neurons, such as olfactory receptor genes. We thus generated our own cell-type specific bulk RNAseq dataset upon FACS sorting genetically marked L2 or L3 neurons. We specifically concentrated our analysis on receptors and ion channels, as these are the most likely candidates to alter the dendritic responses in a cell-type specific manner. From the differentially expressed genes (DEGs) in L3 vs L2, we selected all genes annotated as neurotransmitter receptors (GO:003059, neurotransmitter receptor activity) or ion channels (GO:0005244, voltage-gated monoatomic ion channel activity). We found 34 receptor and ion channel genes that are strongly differentially expressed between L2 and L3 ( |log_2_ fold change| ≥ 3, adjusted p-value < 0.05) (**Fig. 4A**). This list included voltage-gated potassium channels, and chloride channels (**Fig. 4B**). Intriguingly, we found that the second histamine-gated chloride channel, HisCl1 was expressed in lamina neurons and differentially expressed between L2 and L3 (**Fig. 4A,B**). HisCl1 is mostly known to be expressed in glial cells, and more weakly in photoreceptors ^22,24^. Another RNAseq dataset confirms the weak expression in L2 that we found here (Davis et al., 2020) (**Fig. S2A**). HisCl1 can form heteropentameric channels with Ort; however, unlike Ort, HisCl1 exhibits a higher histamine sensitivity and a lower ion conductance ^24,41^, arguing that it could differentially shape a postsynaptic response to histamine.

**Figure 4.**
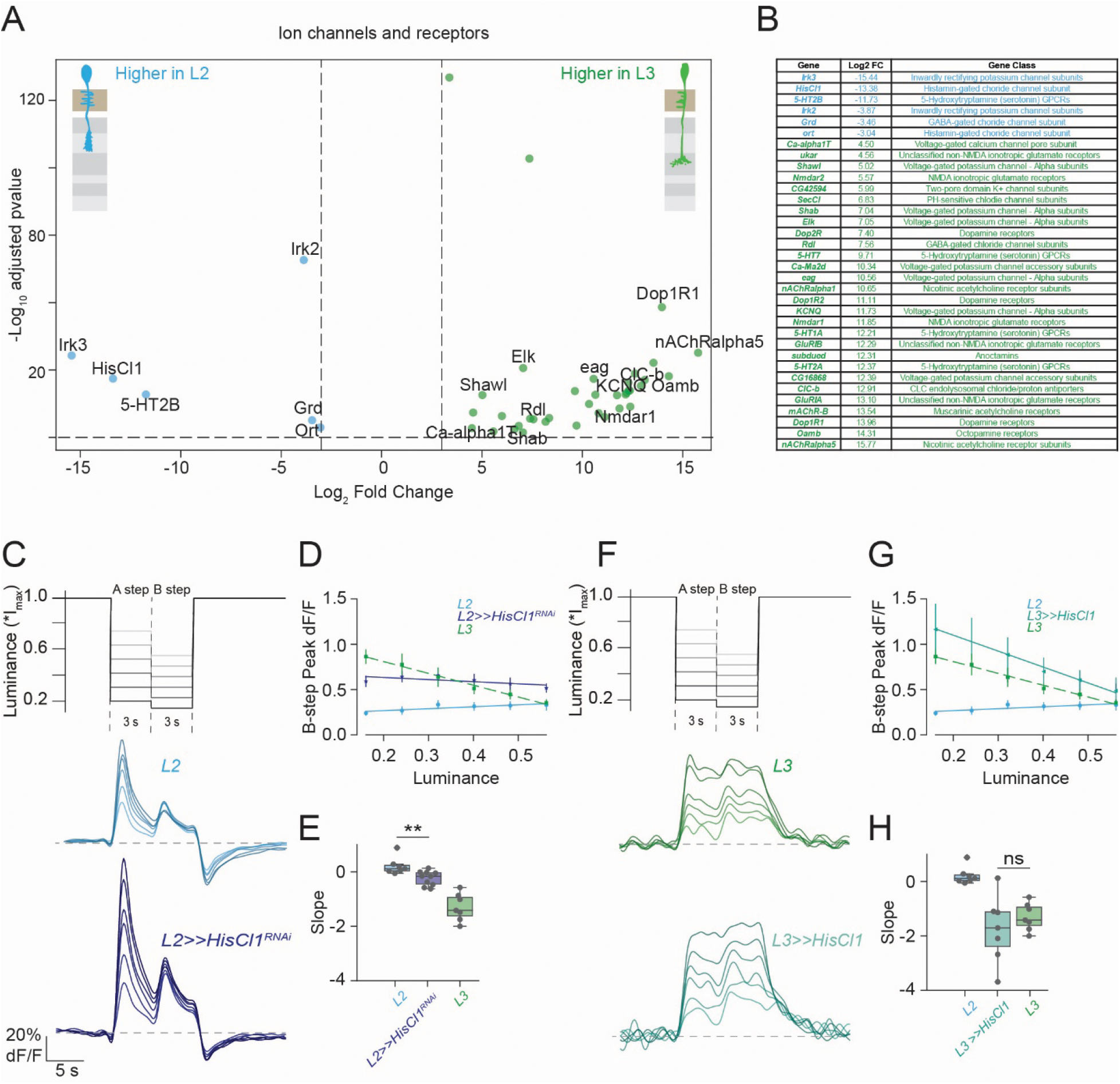
There are many differentially regulated ion channels and receptors between L3 and L2, including the histamine-gated chloride channel HisCl1. **(A)** Volcano plot displaying differentially expressed genes between L3 and L2 neurons. Significance thresholds were set at adjusted p-value < 0.05 and log_2_ fold change ≥ 3 or ≤ -3. Ion channels and receptors enriched in L2 are shown on the left (negative fold change, blue) and those enriched in L3 are shown on the right (positive fold change, green). **(B)** A table showing ion channels and receptors along with their corresponding log_2_ fold change values and gene class. **(C)** L2 responses to the luminance-contrast discrimination stimulus. Two consecutive 3s OFF steps follow an adapting bright period that varied in both luminance and contrast (A step) or varied in luminance with a fixed 25% contrast (B step). Average calcium signals (GCaMP6f) from axon terminals of L2, *L2>>HisCl1^RNAi^* to the luminance contrast discrimination stimulus. **(D)** Quantified mean peak ± SEM (top) responses to B steps plotted as a function of luminance. **(E)** Slopes from linear fits to the data in (D). **(F-H)** Same as (**C-E**) but for L3, *L3>>HisCl1*. **p<0.01, Mann-Whitney test, two-tailed (**E**), ns, not significant. Sample size, L2: n=7 (66), *L2>>HisCl1^RNAi^*: n=10 (86), L3: n=7 (36) and *L3>>HisCl1*: n=7 (84) for (**C-H**).

In the ort-rescue experiment shown above (**Fig. 3**), we found that *ort* mutants still show some responses to visual stimuli (**Fig. 3A**, **Fig. S2B**), further supporting the idea that HisCl1 can contribute to L2 responses. While L2 responses in *ort* mutants appeared more sustained, the low response amplitude did not allow to properly test for contrast and luminance sensitivity (**Fig. S2B**). To test if HisCl1 has a functional role in establishing specific lamina neuron properties, we performed a cell-type specific knockdown of *HisCl1* using RNAi. We tested L2’s visual responses using the luminance-contrast discrimination stimulus and found that L2 neurons upon HisCl1 knockdown responded to OFF steps transiently and with slightly increased amplitudes as compared to the controls (**Fig. 4C**). Importantly, analyzing the B step responses as a function of luminance revealed that the L2 responses upon *HisCl1* knockdown showed a slight dependence on luminance (**Fig. 4D**). Quantification of this luminance dependence as the slope of a linear fit to the response across luminances showed that control responses were close to zero and had a slightly negative slope for *L2>>HisCl1^RNAi^* (**Fig. 4E**). However, the luminance sensitivity upon *HisCl1* knockdown was by far not as strong as those of L3 neurons (**Fig. 4D-E**). To instead ask if HisCl1 is sufficient to confer L2-like properties to L3, we next overexpressed HisCl1 in L3 neurons (**Fig. 4F-H**). L3 neurons overexpressing HisCl1 showed luminance-sensitive, sustained responses that were indistinguishable from control L3 responses (**Fig. 4F-H**). These results suggest that HisCl1 act as a second histamine receptor in lamina neurons where it contributes weakly to shaping L2 responses, but that other factors must play a more prominent role in shaping L2 contrast sensitivity versus L3 luminance sensitivity.

### Many individual candidate genes differentially expressed between L2 and L3 are neither necessary nor sufficient to alter lamina neuron responses

Many receptors and ion channels other than HisCl1 are differentially regulated across L3 vs L2. To determine whether these genes shape the distinct physiological properties of L2 and L3, we systematically manipulated a subset of these candidates. The inward-rectifier potassium channels Irk2 and Irk3 emerged as strong L2-enriched candidates (**Fig. 4A**). Irk2 has been proposed to modulate membrane excitability (Chen et al., 2018; Piermarini et al., 2022), while the less characterized Irk3 is thought to play a role in trafficking or stabilizing other Irk subunits at the plasma membrane (Piermarini et al., 2022). To test whether these channels modulate contrast versus luminance sensitivity, we knocked them down individually in L2. Upon either *Irk2* or *Irk3* knockdown, L2 still showed transient responses to the luminance-contrast discrimination stimulus, which were encoding contrast, as demonstrated by similar responses to the different -25% contrast ‘B steps’ across luminances (**Fig. 5A-C**). Overexpression of Irk2 or Irk3 in L3 maintained normal sustained, luminance-sensitive L3 responses, as B step responses varied across luminances (**Fig. 5D-F**). Because Irk2 and Irk3 are structurally similar and can form both homo- and heterotetrameric channels ^42,43^, we overexpressed a dominant-negative Irk2 isoform (Irk2-AAA) that disrupts Irk2 and Irk3 channels simultaneously ^43^. While the expression of Irk2-AAA in L2 increased calcium signal amplitudes, the responses remained transient and contrast sensitive, as in control L2 neurons (**Fig. 5A-C).**

**Figure 5.**
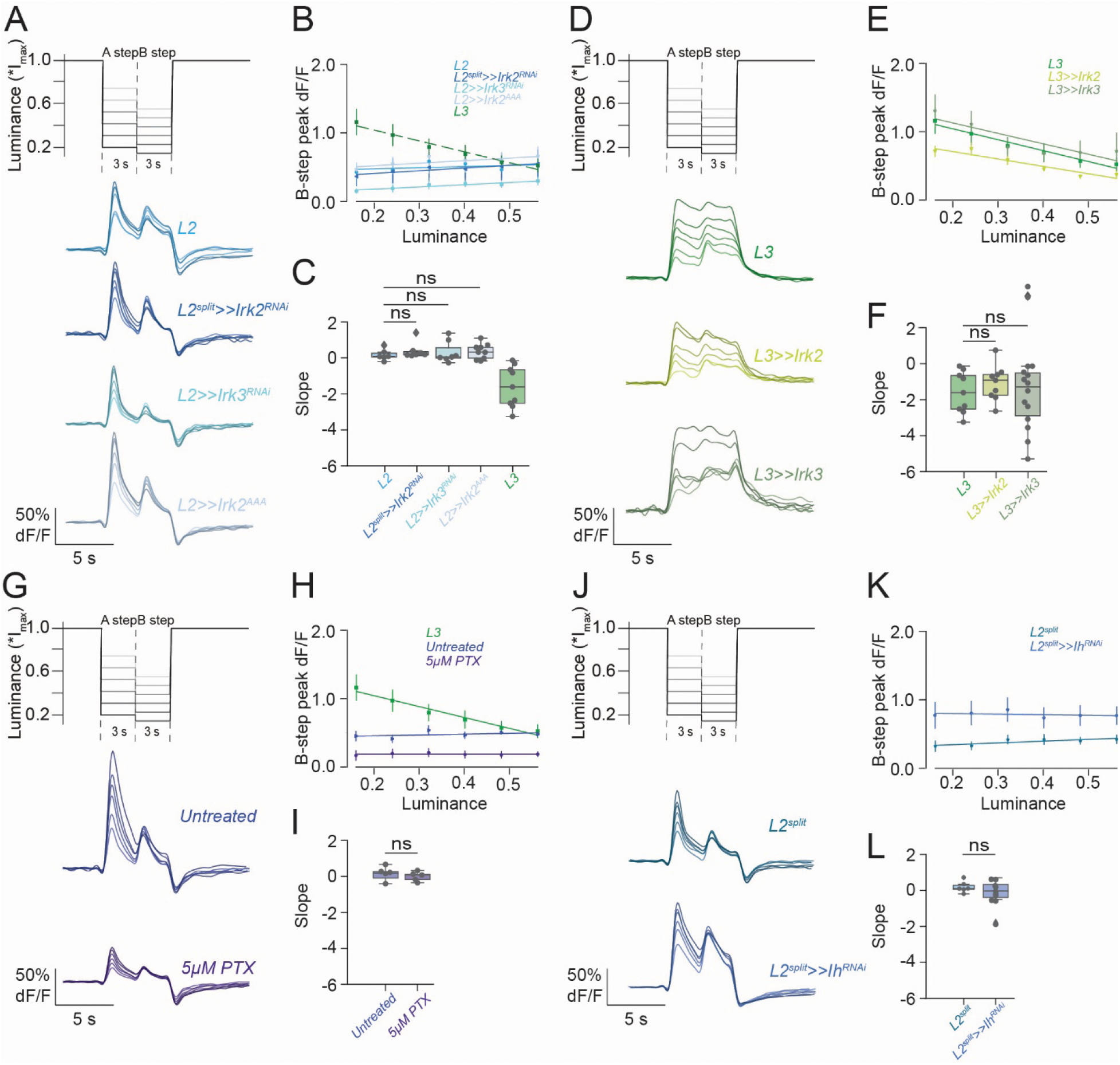
Manipulation of individual L2 enriched ion channels and receptors does not change L2 contrast and L3 luminance sensitivity. **(A).** Average calcium (GCaMP6f) signals from axon terminals of *L2* (blue)*, L2^spli^t>>Irk2^RNAi^, L2>>,Irk3^RNAi^, L2>>Irk2^AAA^* to the luminance-contrast discrimination stimulus. (**B**) Quantified mean peak ± SEM responses to B steps plotted as a function of luminance. (**C**) Slopes from linear fit to the responses in (B). **(D-E)** Same as (**A-C**) but for *L3, L3>>Irk2,* and *L3>>Irk3*. **(G-I)** Same as (**A-C**) but for untreated L2 and 5µM PTX-treated L2. **(J-L)** Same as (**A-C**) but for *L2^split^* control neurons and *L2^split^>>Ih^RNAi^*. ns, not significant. Sample size, *L2*: n=6 (66)*, L2^split^>>Irk2^RNAi^*: n=7 (82)*, L2>>Irk3^RNAi^*: n=7 (53), *L2>>Irk2^AAA^*: n=9 (78)*, L3: n*=9 (56)*, L3>>Irk2*: n=9 (79)*, L3>>Irk3*: n=14 (120)*, Untreated L2*: n=5 (34)*, 5µM PTX Treated L2*: n=5 (25)*, L2^split^>>GcaMP6f*: n=7 (85)*, L2^split^>>Ih^RNAi^*: n=10 (120). Sample sizes are given as number of flies (number of cells).

Another L2-enriched gene, Grd (**Fig. 4A**), encodes an ionotropic GABA_A_ receptor subunit. Using a pharmacological approach, we broadly blocked GABAergic inhibition using picrotoxin (PTX). Upon bath application of 5µM PTX, L2 calcium signals were reduced in amplitude, but still responded transiently to the luminance-contrast discrimination stimulus, and their B step responses did not change with luminance (**Fig. 5 G-I**). Furthermore, a recent modeling study (Borst, 2025) proposed that the hyperpolarization-activated cyclic nucleotide-gated (HCN) channel Ih, which was reported to be present in transient neurons like L1 and L2 but absent in sustained neurons like L3 ^40^, can explain the transient response dynamics in L2. We tested this idea by knocking down *Ih* in L2 neurons. The L2 neurons responded transiently to the luminance-contrast discrimination stimulus upon *Ih* knockdown, and the B step responses did not show luminance dependence and were similar to control L2s (**Fig. 5J-L**). When we further used the stimulus with randomly varying luminance, there were subtle changes in the luminance dependence of L2, such that Ih^dsRNA^ expressing L2 neurons appeared to show slower decay as compared to controls (**Fig. S3A**). However, quantification of the luminance dependence across all flies recorded did not show significant differences between the knockdown and control (**Fig. S3A-D)**.

Turning to the L3-enriched genes (**Fig. 4A**), we investigated the delayed rectifier potassium channel Shab. L3-type neurons in the blowfly visual system exhibit delayed rectifier K+ currents ^26,27^, and Shab is necessary for maintaining slow, sustained potassium (IK) currents in *Drosophila* embryonic neuronal cultures (Peng et al., 2007). Knocking it down in L3 did not alter its characteristic luminance sensitivity (**Fig. S3E-G**). Overexpressing *Shab* in L2 neurons altered its responses amplitude and responses were so low that it was difficult to assess kinetics (**Fig. S3K-N**). We similarly tested the L3-enriched glutamate receptor NMDAR1, and found it to be neither necessary nor sufficient to shape the contrast vs luminance sensitive properties (**Fig. S5E-J**). These results show that none of the differentially regulated channels or receptors tested here were alone required for the core contrast- vs luminance-sensitivity differences between L2 and L3 neurons.

### The L3-specific transcription factor dFezf regulates L3 physiological properties

Testing many individual candidate genes, encoding channels or receptors that are either enriched in L2 or L3, did not alter the physiological response profile of either cell type substantially. While it is plausible that we simply failed to identify the one relevant factor, another possibility is that the system is degenerate, meaning that different genes jointly contribute to the same function. Especially ion channel degeneracy is well established in many different system, from the crab stomatogastric ganglion to hippocampal cell types ^30,32,34,44^. If it is true that a degenerate set of candidate genes establishes L2 vs. L3 properties, it will not be possible to identify the responsible component by manipulating individual factors, and manipulating several candidates simultaneously is experimentally demanding. We hypothesized that, if many different factors jointly contribute to setting up cell-type specific dynamics, these factors could be jointly controlled by an overarching regulator.

We explored this hypothesis by asking if we can find individual factors controlling the joint expression of several channels and/or receptors that are then important for function. Importantly, we had access to candidates, as each lamina neuron type has been reported to express a unique set of transcription factors ^45^. While some transcription factors are known to fully change cell type identity in the lamina ^46,47^, dFezf (also called Earmuff, Erm) is an L3-specific transcription factor that is reported to have a role in the axonal guidance of L3 ^48,49^. In a dFezf mutant, L3 axons mistarget to a more proximal domain of the developing medulla, leading L3 neurons to mistarget to more distal medulla layers. However, L3 dendrites maintain their characteristic L3-like shape and *dFezf* mutant L3 neurons do not turn on other cell-type markers, arguing against a cell-fate change ^48^. We thus tested whether, in addition to its role in axonal targeting, dFezf also has a role in modulating L3 physiological properties. Because *dFezf* mutants are lethal, we used MARCM (Mosaic analysis with a repressible cell marker), a genetic technique to generate homozygous mutant cells in an otherwise heterozygous background, while dominantly labeling these cells with a marker. To be able to record the visual response properties of wild type and *dFezf* mutant L3 clonal cells, we used GCaMP6f as a dominant marker. Lamina specific clone generation (using 27G05-Flp) labelled individual L3 neurons in control and *dFezf mutant* conditions (**Fig. 6A-D**). To distinguish between L2 and L3-like physiological properties, we next used the luminance-contrast discrimination stimulus. Strikingly, the *dFezf mutant* L3 neurons showed transient responses, whereas responses of wild type L3 neurons were sustained (**Fig. 6E**). Control L3 neurons responded similarly to close-by luminances regardless of contrast (regardless of whether this was the A- or B-step), whereas the responses to the six 25% contrast B steps differed (**Fig. 6F,G**), demonstrating luminance sensitivity. *dFezf mutant* L3 neurons respond similarly to the same contrast at varying luminance (**Fig. 6H,I**), implying that L3 neurons lost their luminance information and instead responded like contrast-sensitive L2 neurons. This shows that the dFezf transcription factor is alone required for L3’s luminance sensitivity. To further test the luminance encoding, we recorded calcium responses in the MARCM clones when flies were shown a stimulus of randomly varying luminance flashes. The control L3 neurons showed sustained responses to the OFF steps which did not return to a fixed baseline (**Fig. S4A**). Instead, the *dFezf mutant* L3 neurons responded transiently to the OFF steps before returning to a baseline, similar to previously reported L2 responses (**Fig. S4B**). Quantifying the sustained plateau component of these responses showed that *dFezf mutant* L3 lacked the luminance-encoding sustained component (**Fig. S4C**).

**Figure 6.**
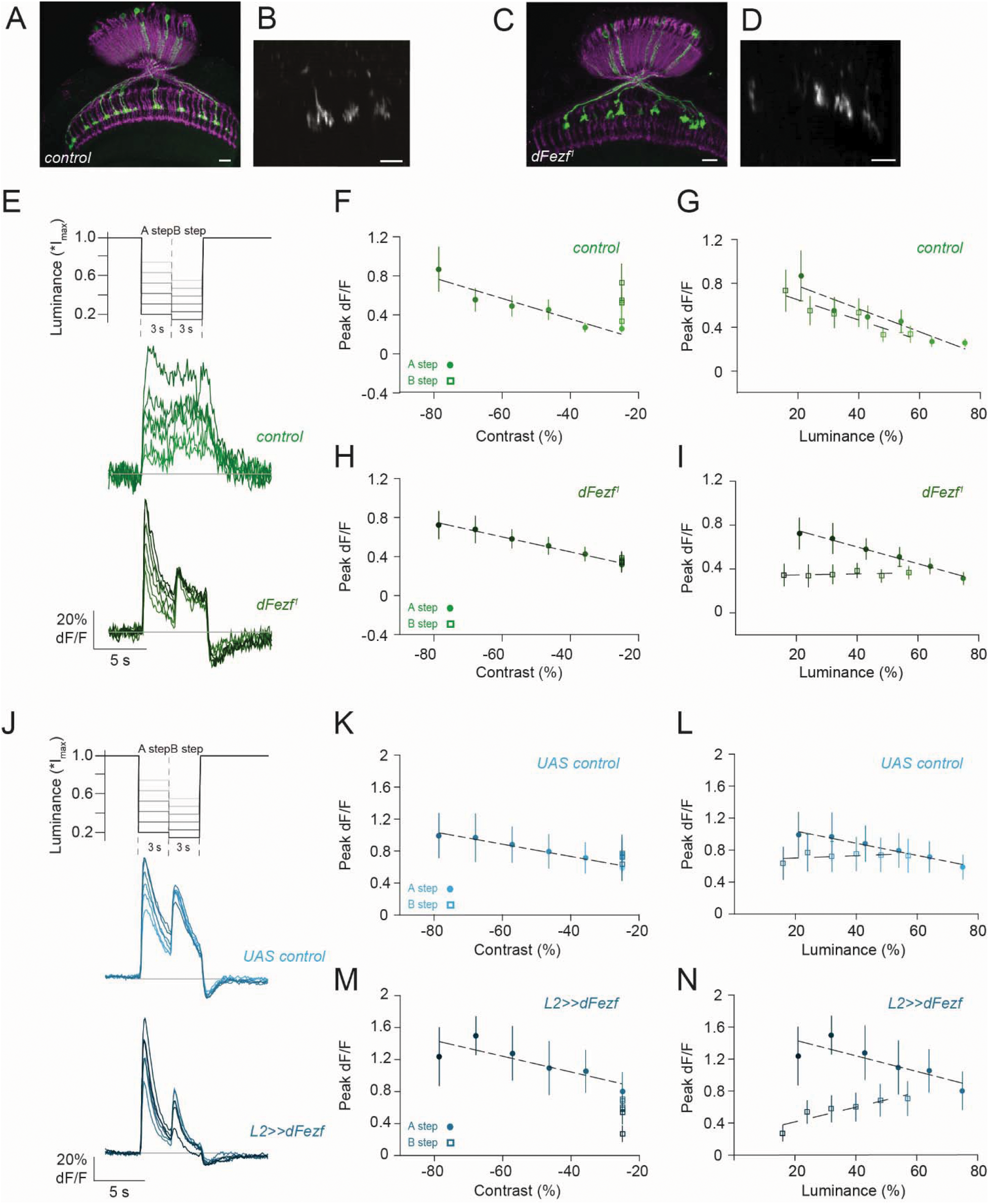
Luminance sensitivity of L3 is established by dFezf. **(A-D)** Control (A, B) or *dFezf*-mutant (C, D) L3 MARCM clones dominantly marked by the expression of GCaMP6f. **(A, C)** Representative confocal image from wild-type (**A**) or *dFezf*-mutants (**C**) visual systems, stained with anti-GFP and 24B10. **(B, D)** Representative in vivo calcium image of wild-type (**B**) or *dFezf*-mutant (**D**) L3 neurons expressing GCaMP6f. Scale bars are 10 μm. **(E)** Average calcium (GCaMP6f) signals from axon terminals of control L3 (green) and *dFezf* mutant L3 (dark green) MARCM clones to the luminance contrast discrimination stimulus. **(F, G)** Peak responses (mean ± SEM) to the two OFF steps plotted as a function of contrast (**F**) and luminance (**G**) for control L3. **(H, I)** Same as (**F, G**) for *dFezf* mutant L3 MARCM clones. **(J-N)** Same as (**E-I**) but for UAS control L2 (blue) or *L2>>dFezf* (dark blue). Sample size, Control L3: n=9 (24), *dFezf* mutant L3: n=12 (47) for (**E-I**), UAS control L2: n=7 (72), *L2>>dFezf*: n=6 (64) for (**J-N**). Sample sizes are given as number of flies (number of cells).

We next asked if dFezf is also sufficient to confer L3-like properties to L2. However, overexpressing dFezf did not change the contrast vs. luminance encoding properties of L2 (**Fig. 6J-N, S4D-H**). This differs from dFezf’s role in axon targeting, where overexpression of dFezf in L2 is sufficient to redirect L2 axon terminals, a phenotype that we confirmed in our overexpression experiments (**Fig. S6 D,E**) (Peng et al., 2018). In summary, these results establish that the transcription factor dFezf is necessary but not sufficient for luminance sensitivity in L3 neurons.

### dFezf jointly regulates many candidate receptors and channels that are differentially regulated between L2 and L3

So far, we have shown that the transcription factor dFezf is required for L3-like physiological responses, but that the same does not hold for the many channels and receptors that we tested. This argues for a degenerate system in which dFezf is jointly controlling the expression of many genes that together are important for L3-specific function. To test this possibility, we next investigated L3-specifc gene expression in the dFezf mutant. Specifically, we FACS sorted L3 neurons from fly brains, in which either only control or *dFezf* mutant L3 neurons were dominantly marked with GCaMP6f, using the same genotypes as for the functional analysis above, and acquired their transcriptomes. Many of the genes that we initially isolated to be strong candidates to confer distinct physiological properties to L2 and L3, namely differentially expressed channels or receptors, were regulated by dFezf (**Fig. 7A,B**). We found that essentially all genes which we tested functionally (in Fig. 4, 5, Fig. S3A, some not shown) depended on dFezf, including e.g. the L2-enriched targets HisCl1, Irk2 and Irk3. These genes were upregulated in a *dFezf* mutant (**Fig. 7B**). Furthermore, many genes which we initially found to be enriched in L3, were also regulated by dFezf, arguing that they are normally activated by dFezf in L3. This includes the potassium channel Shab, and the NMDA receptor. (**Fig. 7B)**. Expanding on the idea that dFezf is activating a host of genes that could jointly, or redundantly control L3 physiological function, we plotted all neurotransmitter receptors, ion channels, and ion channel regulators, which were regulated by dFezf (adj. p-value < 0.05, |log2 fold change| ≥ 0.6 for both, L2 vs. L3, and L3 control MARCM vs L3 *dFezf^1^*MARCM). This list includes many of the genes that we already tested, unexplored CGs, or receptors for serotonin (5-HT1B, 5-HT7) and octopamine (Oamb) neuromodulators (Fig. 7C), of which serotonin has been shown to gain modulate lamina neuron properties without dramatically changing the kinetics ^50^. A consistent picture emerges in which “functional neuronal” genes that are normally higher expressed in L3 are all downregulated in *dFezf*-mutant L3 neurons, and those genes that are normally higher expressed in L2 are upregulated in *dFezf*-mutant L3 neurons (**Fig. 7C)**. Hence, dFezf acts as a transcriptional activator of L3-specific genes and as a transcriptional repressor of L2-specific genes that can regulate neuronal function in L3. Last, we also asked if dFezf overexpression in L2 neurons can turn on L3-specific genes and analyzed L2 cell-type specific RNAseq data with and without overexpression of dFezf. Here, Fezf was not sufficient to turn on the expression of L3-specific genes in L2, and was only able to suppress the expression of some genes that are normally L2-specific. These results are largely in line with our functional analysis, which showed that overexpression of dFezf in L2 did not change L2 physiology (**Fig. 6**). Thus, in line with its adult function, dFezf is necessary but not sufficient to change gene expression of a large set of candidate genes that are well suited to establish L3-like physiological properties.

**Figure 7.**
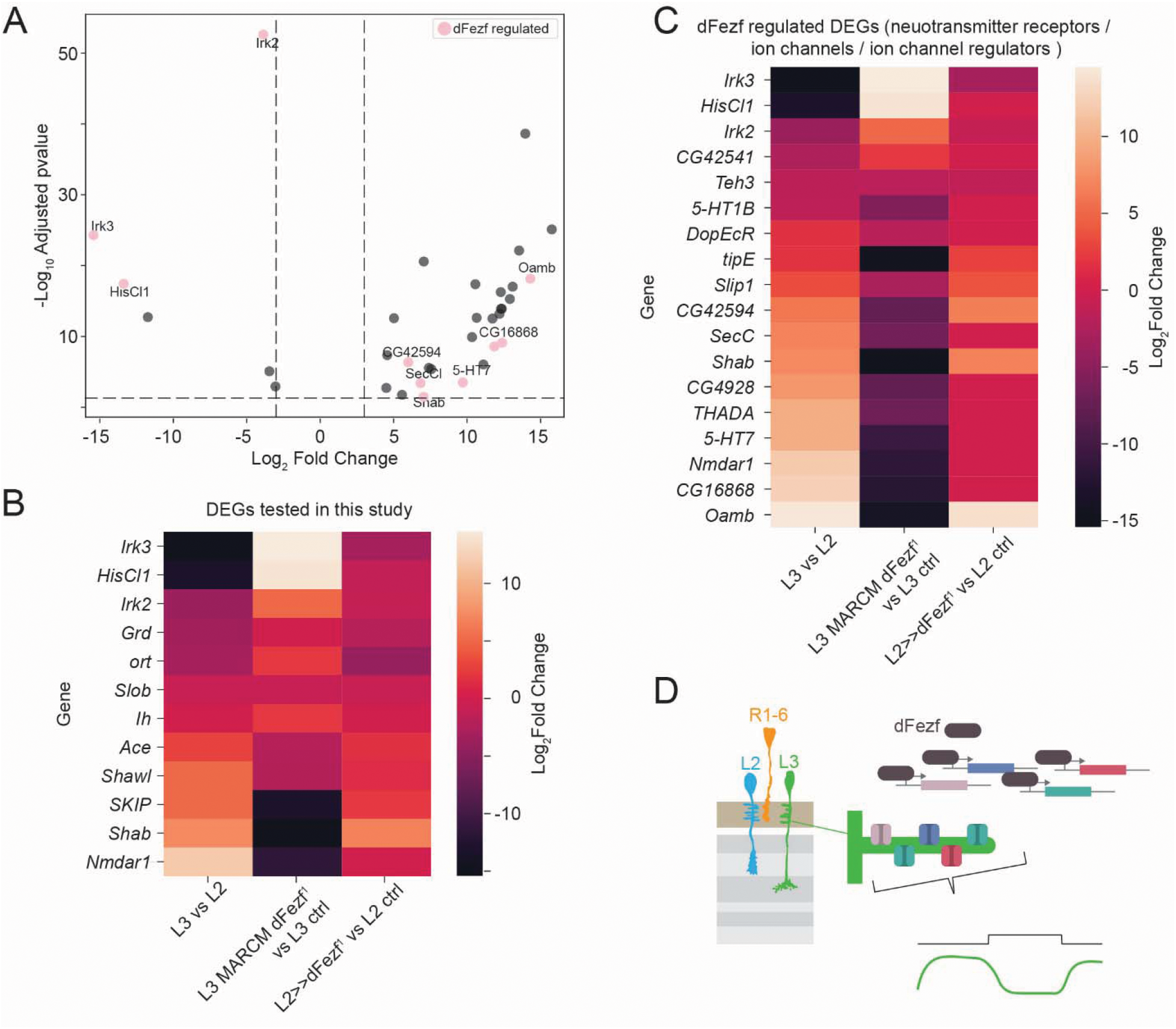
dFezf controls many receptors and channels differentially expressed between L2 and L3. **(A)** Volcano plot displaying differentially expressed genes between L3 and L2 neurons. Significance thresholds were set at adjusted p-value < 0.05 and log_2_ fold change ≥ 3 or ≤ -3. Ion channels and receptors enriched in L2 are shown on the left (negative fold change) and those enriched in L3 are shown on the right (positive fold change). Dfezf regulated genes are highlighted in pink. **(B)** Heatmap showing log_2_ fold change for differential regulated ion channels and receptors in L3 vs L2 neurons that were functionally tested by us (top is high in L2, bottom is high in L3). The middle column shows if these genes are differentially regulated in *dFezf* mutant L3 MARCM clones as compared to controls (top is high in *dFezf* mutant L3 MARCM clones and bottom is high incontrol L3 MARCM clones). The right column shows if these genes are differentially regulated in L2 with and without overexpression of *dFezf* in L2 (top is high in control L2 and bottom is high in *L2>>dFezf*). **(C)** Heatmap showing log_2_ fold change for ion channels (GO:0005244), neurotransmitter receptors (GO:0030594) and ion channel regulator activity (GO:0099106) in L3 vs L2. Significance thresholds were set at adjusted p-value < 0.05 and log_2_ fold change ≥ 0.6 or ≤ -0.6. Columns show the same genotypes as in (B). **(D)** Summary schematics: the L3-specific dFezf transcription is regulated a host of ion channels and receptors, leading to L3-specific physiological responses.

## Discussion

Here, we show how lamina neurons L2 and L3 obtain their distinct cell-type-specific functions, which are important for their specific roles in processing behaviorally relevant information. Despite relying on the same photoreceptor input, L2 versus L3 properties are different and largely independent of circuit interactions downstream of photoreceptors. Cell-type-specific transcriptional profiling identifies compelling candidates to confer distinct neuronal properties to L2 and L3, such as the histamine-gated chloride channel HisCl1, or different potassium channels, but loss of any individual component leads to no or only weak phenotypes. In stark contrast to the lack of strong phenotypes upon silencing individual candidate receptor or ion channel, loss of the L3-specific transcription factor dFezf fully converted L3 luminance sensitivity into the contrast selectivity characteristic of L2 neurons. DFezf in turn controls a range of downstream genes, which include many of the above-mentioned L3- or L2-specific membrane proteins potentially important for function. This argues for a degenerate system, where a single transcription factor regulates a wide repertoire of downstream genes that together jointly regulate cell-type-specific temporal properties.

Our work elucidates how lamina neurons obtain fundamentally different temporal properties starting from the same photoreceptor input, allowing them to encode different basic features of the visual scene. First, our work largely excludes the possibility that a contribution from the green-sensitive R8 (or even the UV-sensitive R7) sets up the distinct visual response properties of lamina neurons. Thus, despite functional evidence that achromatic and chromatic pathways interact ^51–53^, these interactions do not influence the specific lamina neurons properties under investigation here. Second, *ort*-rescue experiments revealed a circuit contribution to the elimination of a luminance sensitive baseline in L2. We did not further investigate the potential cellular substrate for this baseline subtraction, but the publication of recent connectomes offers an opportunity to identify candidate cell types. Elimination of the baseline, which could either be subtractive or divisive, could for example be implemented by inhibitory, glutamatergic or GABAergic, neurons synapsing onto L2. Among the major inputs to L2 are the putative glutamatergic neurons Dm15, Lawf1, the GABAergic feedback neurons C2 and C3, and the putative GABAergic neurons Dm6, Dm1, or Dm17 ^54,55^. Although the specific role for L2 properties has not been tested, C2 tunes the temporal tuning of another postsynaptic neuron, Mi1 ^56^, and Dm neurons are proposed to play a role in input normalization ^18,57^, arguing that they might also act upon lamina targets. Upon excluding differential photoreceptor and strong circuit contributions, we focused on the cell-autonomous mechanisms that differentiate L2 and L3 responses. The idea that lamina neurons properties are regulated by cell-intrinsic mechanisms is also supported by recent work showing that recurrent feedback can sharpen L2 filters but does not change their overall biphasic properties ^29^. Similarly, the dynamics of downstream medulla neurons could be altered by changing cell-type specific gene expression ^58^. We ultimately confirmed the role of cell-autonomous mechanisms by demonstrating a full conversion of L3-like into L2-like properties upon the loss of the dFezf transcription factor from an individual L3 cell – a genetic condition where the rest of the circuit is wild type.

We identified dFezf as a transcription factor that is required to establish L3-like properties. Temporal transcription-factor cascades are a widespread mechanism for generating neuronal diversity, from the diverse cell classes of the *Drosophila* visual system ^59^ to within class-diversity. In the lamina, the homeodomain transcription factor Scarecrow (Scro) establishes “class identity” and specifies lamina neurons L1-L5. Scro then activate other subtype-specific homeodomain transcription factors that specify lamina subtype fates. For the cell types investigated here, Dve (Defective proventriculus) is expressed in L2 neurons, whereas Zfh1 is expressed in L3 ^46,47^. Loss of Dve fully converts the cell type identity of L2 neurons into that of L3 neurons, such that they take on morphological, molecular, and functional characteristics ^47^. This is different to what we find for dFezf in L3. While dFezf has been known as an L3-specific transcription factor, and loss of *dFezf* leads to mistargeting of L3 neurons from the M3 to the M2 layer in the medulla, which is the normal L2-target layer, dFezf does not turn on L2-specific genes ^60^. Furthermore, L3 neurons show cell-type specific morphological features such as a uniquely oriented dendritic tree. This dendritic morphology is maintained in *dFezf* mutant L3 neurons, arguing that its input connectivity is preserved. Last, targeting phenotypes and physiological phenotypes are separable: Overexpression of dFezf in L2 is sufficient to mistarget L2 neurons to the M3 layers, but is not sufficient to induce L3-like visual response properties to these neurons, nor is it sufficient to turn on expression of L3-specific functional neuronal genes. Jointly, these findings argue that dFezf is an L3-specific transcription factor that acts after cell fate has been established to define other cell-type-specific properties: it first acts during early pupal development to establish proper L3 axon targeting and then later, to define L3’s luminance sensitive function in the adult fly visual system. More generally, our work argues that cell-type-specific transcription factors, which have been identified for many cell types, could serve as windows into understanding cell-type specific physiological specializations more broadly across cell types.

To link the dFezf-mediated control of gene expression in the nucleus to L3 physiological properties that are established in the dendrites, we concentrated on receptors and channels as the neuronal cellular substrate that can define dendritic responses. We found that dFezf controls the expression of genes encoding for receptors and channels. Strikingly, the list of genes regulated by dFezf overlaps to a large extend with the receptors and genes that we found to be differentially expressed between L2 and L3. This list included compelling candidates, such as the ‘second’ histamine-gated chloride channel HisCl1, a receptor that could directly influence how lamina neurons respond to their histaminergic photoreceptor input. The other histamine receptor, Ort, is generally thought to be the only one that mediates lamina neuron responses to photoreceptor inputs, but several lines of evidence indicate that this might not be the full story. For example, some lamina neuron responses are retained in *ort* null mutants, and different transcriptome studies largely agree that HisCl1 is weakly expressed in L2, and shows stronger expression as compared to L3 ^40^. While expression is weak, even incorporation of one HisCl1 subunit into a heteropentameric channel that is otherwise mostly composed of Ort subunits could alter postsynaptic properties. Heteropentameric channels composed of HisCl1 and Ort subunits exhibit greater histamine sensitivity but lower single-channel conductance compared to homopentameric Ort channels ^24^, a change which may tune responses to histamine in neurons expressing both genes, even at different stoichiometries. However, functional analysis could only identify a minor contribution to the cell-type specific properties of L2. Thus, although different receptor expression might be the easiest way to diverge different neuronal properties from a common presynaptic input, different or additional mechanisms must be in place.

Combining the requirement of dFezf for L3-specific function, and its ability to jointly regulate many L3-specific genes hence argues for a degenerate system, in which different combination of receptors and/or channels jointly establish L3 function. Ion channel degeneracy has arguably been best studied in the crab stomatogastric ganglion, where different sets of ion channels can establish the same firing pattern of identified cell types ^30,44^. Interestingly, work in this system has further revealed that ion channel expression is correlated, and that homeostatic tuning rules can explain these correlations ^35,61^. Theoretical work has shown that regulating a set of conductances via a single master regulator is advantageous, preventing that a system that fails to control several target functions simultaneously runs out of bounds ^61^. Our data suggest that coordinated transcriptional regulation by a common, yet cell-type specific transcription factor can be one way to achieve functionally meaningful correlated expression. In that scenario, control of ion channel and receptor expression by a common genetic regulatory mechanism is not a byproduct that leads to these gene expression correlations, but instead is the mechanism by which meaningful correlations are established in a cell-type specific manner.

The idea that having a single global regulator of gene expression for controlling downstream precursors of membrane conductances is advantageous ^61^ also directly addresses the question why cell-type specific gene expression control relevant for function is implemented by a single factor, whereas having multiple solution exist at the receptor and channel level. Degenerate systems for controlling neuronal functional properties are robust to perturbations, as also shown here by the lack of strong phenotypes when manipulating individual components. In general, they are also more versatile ^30,31^ and this versatility is easily exploited at the neuronal membrane, where an L3 neuron receives input from many different cell types, including feedforward, feedback and neuromodulatory connections ^54,55^. Our transcriptome dataset includes receptors for major neurotransmitters (e.g., GABA) and neuromodulators (e.g., serotonine and octopamine), suggesting that versatility in transmembrane protein expression can accommodate different functional demands of the system that remain unexplored in this work. We propose that the system that we present here, functionally diverse cell-types at the same level of hierarchy, whose cell-type specific temporal properties have been directly linked to function down to the behavioral level ^10,11,15,17^, will be an excellent system to further understand the trade-offs between ensuring both robustness and versatility in neural systems.

## Resource Availability

All original code will be deposited at Github, and data will be made publicly available via Zenodo upon publication of this study.

## Acknowledgments

We thank Carlotta Martelli and Christopher Schnaitmann for discussion and comments on the manuscript. We are grateful to Simone Renner for excellent technical assistance. This work was supported by the German Research Foundation (DFG) through priority program SPP2205 and has received funding (to MS) by the European Union (ERC, ‘Adaptive Vision’, Grant Agreement No. 101045003). Views and opinions expressed are however those of the author(s) only and do not necessarily reflect those of the European Union or the European Research Council Executive Agency. Neither the European Union nor the granting authority can be held responsible for them.

## Author contributions

Conceptualization: KS, MS

Formal analysis: KS,

NW Software: KS, NW, MI

Methodology: MI, IJS, JA

Visualization: KS, NW, IJS, MS

Funding Acquisition: MS

Supervision: MS, MP

Writing – Original Draft: KS, MS

Writing – Reviewing and editing. NW, KS, MI, JA, MS

## Declaration of generative AI and AI-assisted technologies in the manuscript preparation process

No AI was used for writing the manuscript. ChatGPT and Gemini were used to correct grammar in the methods section, as well as for literature search, and but all original sources were checked by the authors. The authors take full responsibility for the content of the published article.

## Methods

### Key Resources Table

| **Reagent or Resource** | **Source** | **Identifier** |
|---|---|---|
| **Experimental Models: Organisms/Strains** |  |  |
| *D. melanogaster: wt IsoD1* | Gohl et al. 2011 [1] | NA |
| *D. melanogaster: L2<sup>21Dhh</sup>-Gal4* | Rister et al. 2007 [2] | NA |
| *D. melanogaster: L3<sup>MH56</sup>-Gal4:* | Timofeev et al. 2012 [3] | NA |
| *D. melanogaster: L3<sup>0595</sup>-Gal4 (PBac{IT.GAL4}0595)* | Silies et al. 2013 [4] | NA |
| *D. melanogaster: UAS-GCaMP6f.attP40* | Bloomington Drosophila Stock Center (BDSC) (BDSC) | RRID: BDSC_42747 |
| *D. melanogaster: UAS-GCaMP6f.VK00005* | Bloomington Drosophila Stock Center (BDSC) | RRID: BDSC_52869 |
| *D. melanogaster: L2<sup>R16H03</sup>-lexA.attP40* | Bloomington Drosophila Stock Center (BDSC) | RRID: BDSC_52510 |
| *D. melanogaster: L3[9-9]-Gal4* | Nern et al. 2008 [5] | NA |
| *D. melanogaster: 11-164-Gal4* | Nern et al. 2008 [5] | NA |
| *D. melanogaster: LexAop2-GCaMP6f.attP5* | Bloomington Drosophila Stock Center (BDSC) | RRID: BDSC_44277 |
| *D. melanogaster: L2<sup>Split</sup>-Gal4 R53G02-p65.AD.attP40; R29G11-GAL4.DBD.attP2* | Bloomington Drosophila Stock Center (BDSC) | RRID: BDSC_86946 |
| *D. melanogaster: ort<sup>1</sup>* | Bloomington Drosophila Stock Center (BDSC) | RRID: BDSC_1133 |
| *D. melanogaster: Df(3R)BSC809* | Bloomington Drosophila Stock Center (BDSC) | RRID: BDSC_27380 |
| *D. melanogaster: UAS-HisCl1-RNAi P{KK112578}VIE-260B* | Vienna Drosophila Resource Center | VK104966 |
| *D. melanogaster: UAS-Dcr2* | Bloomington Drosophila Stock Center (BDSC) | RRID: BDSC_24651 |
| *D. melanogaster: UAS-hisCl1.attp5* | Schnaitman et al. 2018 [6] | NA |
| *D. melanogaster: lexAop-myr::tdTomato.attP5* | Chen et al. 2014 [7] | NA |
| *D. melanogaster: 27G05-FLP (X)* | Janelia Research Campus | NA |
| *D. melanogaster: 79C23S-RSRT-STOP-RSRTsmFPV5-2A-LexA* | Peng et al. 2018 [8] | NA |
| *D. melanogaster: dFezf<sup>1</sup>* | Weng et al. 2010 [9] | NA |
| *D. melanogaster: UAS-dFezf<sup>1</sup>-3xHA* | Janssens et al. 2014 [10] | NA |
| *D. melanogaster: ninaE-GCaMP6f* | Asteriti et al. 2017 [11] |  |
| *D. melanogaster: norpA<sup>36</sup>* | Bloomquist et al. 1988 [12] | NA |
| *D. melanogaster: rh1-norpA* | Wernet et al. 2012 [13] | NA |
| *D. melanogaster: panR8-norpA* | Wernet et al. 2012 [13] | NA |
| *D. melanogaster: L3<sup>0221</sup>-QF PBac{IS.QF.w−}0211* | Silies Lab | NA |
| *D. melanogaster: L2<sup>19D12</sup>-LexA.attP40* | Bloomington Drosophila Stock Center (BDSC) | RRID: BDSC_ 52545 |
| *D. melanogaster: UAS-Valium10.attP2* | Bloomington Drosophila Stock Center (BDSC) | RRID: BDSC_35786 |
| *D. melanogaster: Irk2-RNAi P{GD203}v4341* | Vienna Drosophila Resource Center | VDRC4341 |
| *D. melanogaster: Irk3-RNAi P{GD1764}v3886* | Vienna Drosophila Resource Center | VDRC3886 |
| *D. melanogaster: UAS-Irk2-AAA* | Dahal et al. 2012 [14] | NA |
| *D. melanogaster: UAS-Irk3-AAA* | Dahal et al. 2012 [14] | NA |
| *D. melanogaster: UAS-Irk2wt* | Dahal et al. 2012 [14] | NA |
| *D. melanogaster: UAS-Irk3wt* | Dahal et al. 2012 [14] | NA |
| *D. melanogaster: NMDAR1-RNAi {TRiP.HMS02200}attP40* | Bloomington Drosophila Stock Center (BDSC) | RRID: BDSC_41667 |
| *D. melanogaster: UAS-NMDAR1.Exel.2* | Bloomington Drosophila Stock Center (BDSC) | RRID: BDSC_8275 |
| *D. melanogaster: Shab-RNAi {TRiP.JF01823}attP2* | Bloomington Drosophila Stock Center (BDSC) | RRID: BDSC_25805 |
| *D. melanogaster: Ih-RNAi {TRiP.HMC03319}attP2* | Bloomington Drosophila Stock Center (BDC) | RRID:BDSC_51765 |
| *D. melanogaster: UAS-Shab.attP40* | Hürkey et al. 2023 [15] | RRID:BDSC_602187 |
| **Antibodies** |  |  |
| α-GFP chicken, polyclonal | Abcam | Cat#13970<br>RRID:AB_300798 |
| α-chaoptin, mouse, polyclonal | DSHB | Cat# 24B10;<br>RRID:AB_528161 |
| Goat anti-chicken IgG (H + L) Alexa Fluor 488 | Thermo Fisher scientific | Cat# A-11039;<br>RRID:AB_142924 |
| Goat anti-mouse IgG (H + L) Alexa Fluor 647 | Thermo Fisher scientific | Cat# A-21236;<br>RRID:AB_141725 |
| **Chemical compound, drug** |  |  |
| Picrotoxin | Sigma Aldrich | RRID:R284556 |
| **Software** |  |  |
| Python | Python Software Foundation | https://www.python.org |
| MATLAB | Mathworks | http://www.mathworks.com |
| ImageJ | National Institutes of Health | https://imagej.nih.gov/ij |

## Experimental Model Details

Flies for experiments were reared on standard molasses food under a 12:12 h light–dark cycle at 25 °C and 60% humidity. Two-photon experiments were conducted at room temperature. All experiments were performed using 5-7 day old, red-eyed female flies lacking balancer chromosomes GCaMP6 expressed cell-type specifically in Rh1, L2, or L3 neurons to record their visual response properties. RNAi and transgenic stocks used in this study were previously published and validated. Genotypes of all *Drosophila* strains used for experiments are listed in the table below. Exact genotypes are given in the table below.

To record from L2 and L3 neurons in *norpA* mutants, we recorded form *norpA^36^* mutant males. To record from *ort* mutants, we recorded from *ort^1^* in trans to the *Df(3R)BSC809* deficiency, which uncovers the *ort^1^*but not the ninaE locus, at which the *ort^1^* chromosome also carries a mutant.

To investigate L3 neurons in *dFezf^1^*null mutants we used the MARCM (mosaic analysis with a repressible cell marker) technique^16^ to generate homozygous mutant clones in an otherwise heterozygous *dFezf^1^*background. Flp recombinase expression was restricted to lamina precursor cells using the 27G05-FLP driver, and an L3-specific 9-9-Gal4 driver ensured expression of an R recombinase in homozygous wildtype (control) or *dFezf^1^* mutant L3 neurons. This in turn resulted in removal of the STOP cassette in the *BRP-RSRT-STOP-RSRT-smGFPV5-2A-LexA* construct, leading to LexA expression, which activated the sustained expression of lexAop-GCaMP6f to mark and visualize the L3 clones.

### Genotypes used in this study

| Name | Genotype | Figure |
|---|---|---|
| *L2* | *w+; UAS-GCaMP6f / +; L2[21Dhh]-Gal4 / +* | Fig. 1 B-E, 2C , G-H, 5 A-C, G-I, S3 H-J |
| *L3* | *w+; L3[MH56]-Gal4 / +; UAS-GCaMP6f / +* | Fig. 1 B-C, F-G, 2C, 4D-G |
| *L2, norpA<sup>36</sup>* | *norpA[36]; lexAop-GCaMP6f / +; L2[19D12]-LexA / +* | Fig. 2 E |
| *L2, control* | *w+; rh1-norpA / lexAop-GCaMP6f; L2[19D12]-LexA / +* | Fig. 2 E,F |
| *L2, Rh1 rescue* | *norpA[36]; rh1-norpA / lexAop-GCaMP6f; L2[19D12]-LexA / +* | Fig. 2 E,F |
| *L3, norpA<sup>36</sup>* | *norpA[36];QUAS-GCaMP6f / +; L3[0221]-QF / +* | Fig. 2 E |
| *L3 control* | *w+; rh1-norpA / QUAS-GCaMP6f; L3[0221]-QF / +* | Fig. 2 E,F |
| *L3, Rh1 rescue* | *norpA[36]; rh1-norpA / QUAS-GCaMP6f; / L3[0221]-QF / +* | Fig. 2 E,F |
| *L2. control* | *w+; panR8-norpA / lexAop-GCaMP6f; L2[19D12]-LexA / +* | Fig. 2 H,I |
| *L2 Rh5+6 rescue* | *norpA[36]; panR8-norpA / lexAop-GCaMP6f; L2[19D12]-LexA / +* | Fig. 2 H,I |
| *L3, control* | *w+; panR8-norpA / QUAS-GCaMP6f; / L3[0221]-QF / +* | Fig. 2 H,I |
| *L3 Rh5+6 rescue* | *norpA[36]; panR8-norpA / QUAS-GCaMP6f; / L3[0221]-QF / +* | Fig. 2 H,I |
| *ninaE-GCaMP6f* | *w+; ninaE-GCaMP6f / +; + / +* | Fig. S1 |
| *L2 ort +/-* | *w+; UAS-ort / UAS-GCaMP6f; L2[21Dhh]-Gal4, ort[1],ninaE[1] / +* | Fig. 3 A,C,E |
| *L2 ort -/-* | *w+; +/ UAS-GCaMP6f; L2[21Dhh]-Gal4, ort[1],ninaE[1] / Df(3R)BSC809* | Fig. 3 A |
| *L2 ort rescue* | *w+; UAS-ort / UAS-GCaMP6f; L2[21Dhh]-Gal4, ort[1],ninaE[1] / Df(3R)BSC809* | Fig. 3 A,C,E |
| *L3 ort +/-* | *w+; UAS-ort / UAS-GCaMP6f; L3[0595]-Gal4, ort[1],ninaE[1] / +* | Fig. 3 B,D,E |
| *L3 ort -/-* | *w+; + / UAS-GCaMP6f; L3[0595]-Gal4, ort[1],ninaE[1] / DfBSC809* | Fig. 3 B |
| *L3 ort rescue* | *w+; UAS-ort / UAS-GCaMP6f; L3[0595]-Gal4, ort[1],ninaE[1] / Df(3R)BSC809* | Fig. 3 B,D,E |
| *L2>>GCaMP6f; ort<sup>1</sup>/+* | *w+; L3[MH56]-Gal4, P{20XUAS-IVS-GCaMP6f}attP40/+; L2[21Dhh]-Gal4, ort1/+* | Fig. S2 B |
| *L2>>GCaMP6f; ort<sup>1</sup>/df* | *w+; L3[MH56]-Gal4, P{20XUAS-IVS-GCaMP6f}attP40/+; L2[21Dhh]-Gal4, ort1/ort(Df)* | Fig. S2 B |
| *L2* | *w+; P{20XUAS-IVS-GCaMP6f}attP40/+; L2[21Dhh]-Gal4/ P{w[+mC]=UAS-Dcr-2.D}10* | Fig. 4 C-E, G,H |
| *L2>>HisCl1<sup>RNAi</sup>* | *w+; P{20XUAS-IVS-GCaMP6f}attP40/ UAS-HisCl1-RNAi; L2[21Dhh]-Gal4/ P{w[+mC]=UAS-Dcr-2.D}10* | Fig. 4 C-E |
| *L3* | *w+; L3[MH56]-Gal4/+; P{20XUAS-IVS-GCaMP6f}VK00005/+* | Fig. 4 D-H |
| *L3>>HisCl1* | *w+; L3[MH56]-Gal4/ UAS-hisCl1(su(Hw)attp5); P{20XUAS-IVS-GCaMP6f}VK00005/+* | Fig. 4 F-H |
| *L3* | *w+; L3[MH56]-Gal4/ +; P{20XUAS-IVS-GCaMP6f}VK00005/P{UAS-GFP.VALIUM10}attP2* | Fig 5 B-F,H, S3 B-G, I-J |
| *L2<sup>split</sup>* | *w+; P{R53G02-p65.AD}attP40/ P{20XUAS-IVS-GCaMP6f}attP40; P{ R29G11-GAL4.DBD}attP2/+* | Fig. 5 J-L, S3 A, C-D, H-N |
| *L2<sup>split</sup>>>Irk2<sup>RNAi</sup>* | *w+; P{R53G02-p65.AD}attP40/ P{20XUAS-IVS-GCaMP6f}attP40; P{ R29G11-GAL4.DBD}attP2/ P{GD203}v4341* | Fig. 5 A-C |
| *L2>>Irk3<sup>RNAi</sup>* | *w+; P{20XUAS-IVS-GCaMP6f}attP40/+; L2[21Dhh]-Gal4/ P{GD1764}v3886* | Fig. 5 A-C |
| *L2>>Irk2<sup>AAA</sup>* | *w+; P{20XUAS-IVS-GCaMP6f}attP40/+; L2[21Dhh]-Gal4/ P{UAS-Irk2A-AAA}13* | Fig. 5 A-C |
| *L2>>NMDAR1* | *w+; P{20XUAS-IVS-GCaMP6f}attP40/ P{UAS-Nmdar1.Exel}2; L2[21Dhh]-Gal4/+* | Fig. S3 H-J |
| *L2<sup>split</sup>>>Shab* | *w+; P{R53G02-p65.AD}attP40/ P{20XUAS-IVS-GCaMP6f}attP40; P{ R29G11-GAL4.DBD}attP2/10xUAS-Shab-pJFRC81attP2* | Fig. S3 K-N |
| *L3>>Irk2* | *w+; L3[MH56]-Gal4/ +; P{20XUAS-IVS-GCaMP6f}VK00005/ P{UAS-Irk2A-wt}2* | Fig. 5 D-F |
| *L3>>Irk3* | *w+; L3[MH56]-Gal4/ +; P{20XUAS-IVS-GCaMP6f}VK00005/ P{UAS-Irk3-wt}1* | Fig. 5 D-F |
| *L2<sup>split</sup>>>Ih<sup>RNAi</sup>* | *w+; P{R53G02-p65.AD}attP40/ P{20XUAS-IVS-GCaMP6f}attP40; P{ R29G11-GAL4.DBD}attP2/P{y[+t7.7] v[+t1.8]=TRiP.HMC03319}attP2* | Fig 5. J-L, S3A, C-D |
| *L3>>NMDAR1<sup>RNAi</sup>* | *w+; L3[MH56]-Gal4/ P{ TRiP.HMS02200}attP40; P{20XUAS-IVS-GCaMP6f}VK00005/+* | Fig. S3 E-G |
| *L3>>dcR2,Shab<sup>RNAi</sup>* | *w+; L3[MH56]-Gal4/ P{20XUAS-IVS-GCaMP6f}attP40; P{w[+mC]=UAS-Dcr-2.D}10/ P{y[+t7.7] v[+t1.8]=TRiP.JF01823}attP2* | Fig. S3 E-G |
| *L3 MARCM control* | *w+; FRT40, lexAop-GCaMP6fp10 / tub-Gal80, FRT40, 27G05-Flp; 9-9-Gal4, UAS-R / BRP-RSRT-STOP-RSRT-smGFPV5-2A-LexA* | Fig. 6 A,B, E-G, S4 A, C, 7 B,C |
| *L3 MARCM dFezf1* | *w+; dFezf[1] FRT40, lexAop-GCaMP6fp10 / tub-Gal80, FRT40, 27G05-Flp; 9-9-Gal4,UAS-R / BRP-RSRT-STOP-RSRT-smGFPV5-2A-LexA* | Fig. 6 C,D, E, H-I S4 B-C, 7 B,C |
| *UAS control* | *w+; lexAop-GCaMP6fp10attp5 / L2[16HO3]-LexA, lexAop-myrtdTOM; + / UAS-dFezf::HA* | Fig. 6 J-L, S4 D,F-H |
| *L2>>dFezf* | *w+; lexAop-GCaMP6fp10attp5 / L2[16HO3]-LexA,lexAop-myrtdTOM; 11-164-Gal4 / UAS-dFezf::HA* | Fig. 6 J,M,N, S4 E,G,H, 7 B,C |
| *Gal4 ctrl* | *w+; lexAop-GCaMP6fp10attp5 / L2[16HO3]-LexA, lexAop-myrtdTOM; 11-164-Gal4 / +* | Fig. 7 B-C |

## Method details

### Two-photon imaging

Two-photon imaging was performed at 20 °C. Flies were cold-anesthetized and mounted on stainless-steel foil with an opening for head and thorax. The head was fixed using UV-curable adhesive (Bondic) and legs immobilized with bee wax. The head was tilted to expose the posterior cuticle, where a small window was opened; excess cuticle and trachea were removed with fine forceps (#5, Biologie/Inox). To perform calcium imaging of the dendrites, the medulla was carefully removed using forceps to expose the underlying lamina, which has the L2 and L3 dendrites. To optimize optical access to the dendrites, the head was adjusted with a slight downward tilt on the targeted side, positioning it at an angle of approximately 30° below the horizontal plane. During imaging, the brain was perfused with a carboxygenated saline solution containing: 103 mM NaCl, 3 mM KCl, 5 mM TES, 1 mM NaH2PO4, 4 mM MgCl2, 1.5 mM CaCl2, 10 mM trehalose, 10 mM glucose, 7 mM sucrose, and 26 mM NaHCO3 (pH ∼ 7.3, bubbled with 95% O2 / 5% CO2.). For dissections, the same solution was used without calcium and sugars. Imaging was performed on a Bruker Ultima microscope (20×/NA1.0 objective, Leica) with 930 nm excitation (5–15 mW at the objective; Toptica) Emission was collected through a SP680 short-pass filter, a 560 lpxr dichroic mirror, and a 525/70 emission filter. Data were acquired using PrairieView software at ∼10 Hz frame rate with an optical zoom of 6x - 8x. Imaging L2/L3 dendrites in the lamina was technically challenging because they are located beneath the retina and partially covered by the medulla. Consequently, detectable dendritic responses were observed in only ∼40-50% of flies, compared to nearly all fly recordings from L2 and L3 axon terminals.

### Pharmacology

Handling, storage, and disposal of all pharmacological compounds strictly adhered to institutional protocols and manufacturer safety data sheets (SDS). Concentrated stocks of Picrotoxin (PTX; Sigma Aldrich, R284556) were prepared in deionized water, aliquoted, and maintained at -20 °C for a maximum of two months. On the day of recording, these stocks were diluted directly into the calcium imaging buffer to yield fresh working solutions. PTX was delivered via bath application to the brain tissue, without any perfusion pre- or post-application. Finally, to allow adequate diffusion and tissue penetration, the preparation was incubated in the toxin for 5 minutes prior to the imaging.

### Visual stimulation

All visual stimuli were generated using custom-written software in either PsychoPy ^17^ or C++ and OpenGL and delivered using LightCrafter projector (Texas Instruments, Dallas, TX, USA) at a frame rate of 100 Hz. The stimuli were filtered through a 482/18 band-pass filter and a ND1.0 neutral density filter and project onto a 9 x 9 cm rear-projection screen placed in front of the fly, spanning 80° of the visual field in both azimuth and elevation. Screen luminance was determined by measuring power at the screen using a digital optical power meter with a 9.5 mm aperture Si photodiode (at 482 nm). The maximum screen luminance I_max_ was ∼1.9∗10^5^ photons∗s^−1^∗receptor^-1^.

### Full-field flashes

The stimulus consisted of periodic, alternating full field ON and OFF flashes covering the entire screen, each lasting 5 s. Each stimulus epoch was repeated at least seven times.

### Full-field flashes from an intermediate gray

The stimulus consisted of random, alternating light and dark flashes of full contrast, lasting 2 s, interleaved with an intermediate gray epoch lasting 4 s. Each stimulus epoch was repeated at least five times.

### Full field flashes of randomly varying luminance stimulus

The stimulus consisted either of 10 s full-field flashes of 5 different intensities (0, 0.25, 0.5, 0.75, 1* of I_max_) (I_max_ =1.87∗10^5^ photons∗s^−1^∗receptor^-1^, Fig. 3, S1, S4) or of 7 s full-field flashes at 11-luminance levels (0, 0.1, 0.2, 0.3, 0.4, 0.5, 0.6, 0.7, 0.8, 0.9, and 1* of I_max_) (I_max_ =1.93∗10^5^ photons∗s^−1^∗receptor^-1^, Fig. 1, S3). Epochs were randomized and repeated at least three times.

### Luminance-contrast discrimination stimulus

To distinguish contrast-sensitive from luminance-sensitive neurons, we used a stimulus protocol adapted from (^18, 19^), which consists of a 30 s adapting bright period followed two consecutive OFF steps (A and B), each lasting 3 s. The first “A step” comprises one of six linearly decreasing luminance values, producing six different luminance and six different Weber contrast values relative to the adapting bright period. The subsequent B step also comprised of one of six decreasing luminance values, chosen based on the preceding A step, resulting in six uniform 25% Weber contrast steps. A-B epoch pairs were randomized and repeated at least three times.

### Visual stimulation at different wavelengths

To assess the spectral efficiency of L2 and L3, we used a multispectral LED-based light synthesizer with a diffraction grating ^20^. The light from a series of LEDs is passed through a diffraction grating to a collimating lens and then into the light fiber, which is projected into the fly’s eye. Stimuli consisted of LEDs spanning UV to red (367, 373, 385, 405, 420, 439, 451, 469, 485, 500, 518, 530, 554, 573, 590, 626, 651, 660, 672, and 682 nm; ∼10 nm bandwidth), calibrated with a spectroradiometer (Flame, Ocean Optics). LED intensities were controlled via 12-bit pulse-width modulation (1 kHz) to generate an isoquantal stimulus set. Visual stimulation was synchronized with photomultiplier tube (PMT) shutters to prevent light contamination. Shutter closing and opening times were 18 ms and 6 ms, respectively, determined from stimulus bleedthrough. The imaging protocol (PrairieView, Mark Points) triggered shutter closure 18 ms before stimulus onset and reopening 6 ms before stimulus offset, ensuring complete shuttering during stimulus presentation. Peak responses to each wavelength were used to construct spectral efficiency curves.

## Quantification and Statistical Analysis

Data processing and analysis for calcium imaging was performed using Matlab (Mathworks, Natick, MA) and Python 2.7. Acquired images were aligned to a reference stack (maximum intensity projection of the first 30 frames) using cross-correlation after Fourier transformation or motion correction based on a hidden Markov model (SIMA Python package). Regions of interest (ROIs) were manually selected. Mean fluorescence per ROI was computed for each frame, background-subtracted to generate a time series. Neural and stimulus signals were interpolated to 10 Hz. Activity was expressed as ΔF/F_0_, with F_0_ defined as the mean fluorescence over the full trace unless mentioned otherwise. For all stimuli except “visual stimulation at different wavelengths,” individual ROI responses were correlated with the stimulus and filtered based on the correlation coefficient; only negatively correlated responses were included in the analysis for L2 and L3 neurons, and only positively correlated responses were for R1-R6 ^21^. Mean responses were first computed across ROIs within each fly and then averaged across flies. Manually selected responsive flies were included for visual stimulation at different wavelengths. Using the Mark Points function in PrairieView, PMT shutters were closed during light presentation, resulting in zero fluorescence. Such stimulus epochs were identified, and epochs longer than 1 s were retained to exclude noise. Traces were truncated at epochs exceeding 4 s to remove post-stimulus activity. Calcium responses to repeated presentations of the same LED were then averaged. All statistical analyses were performed across flies. For pairwise statistical comparisons, normality of the data distributions was evaluated using both the Shapiro-Wilk and Kolmogorov-Smirnov tests (p > 0.05). A two-tailed, unpaired Student’s t-test for normally distributed datasets, whereas the non-parametric Mann-Whitney U (Wilcoxon rank-sum) test for non-normally distributed datasets was used.

### Full-field flashes

Peak and plateau calcium responses were defined as the maximum response within 3 s following the OFF stimulus and mean response during the final 1 s of the OFF stimulus respectively, relative to the mean baseline over the 2 s preceding the step. The percentage of peak retained at plateau was calculated as (plateau/peak) x 100.

### Full-field flashes from an intermediate gray

ΔF/F_0_ was computed using the mean fluorescence during the gray period as baseline (F_0_). Peak OFF responses to gray-to-OFF or ON-to-gray transitions were quantified relative to the mean fluorescence over the 1 s preceding the step. Statistical comparisons between equivalent OFF steps as mentioned in the figure legends.

### Flashes of randomly varying luminance stimulus

For the 10 s version of the stimulus (5 different luminance values), ΔF/F_0_ was computed using the mean fluorescence during the 100% ON step as baseline (F). Plateau responses were defined as the maximal change in calcium signal during the final 2 s of the response relative to the mean fluorescence during the 2 s preceding the stimulus step. Differences between responses to steps ending at the same intensity were assessed using one-way ANOVA.

For the 7 s version of the stimulus (11 different luminance values) plateau responses were computed as the mean ΔF/F_0_ during the final 500 ms of each stimulus epoch. These responses were then related to stimulus luminance by fitting an exponential function of the form F(l)=N_0 *_ e^l/τ^ +c, where F(l) represents the plateau ΔF/F_0_ at luminance l, N_0_ is the response at the lowest luminance, c is a constant offset, and τ is the decay constant reported as the exponent. Fits were performed using bootstrap resampling (5000 iterations) to estimate parameter distributions. The exponential parameter τ quantifies response non-linearity, with τ>0 indicating non-linear increase, τ<0 indicating non-linear decrease, and τ≈0 indicating a linear relationship. Statistical comparisons between exponents as mentioned in the figure legends.

### Luminance-contrast discrimination stimulus

Peak responses were defined as the maximum ΔF/F of each A- and B-step, relative to the mean baseline activity during the final 2 s of the adaptation period. Luminance dependence was quantified by fitting linear regression models to the response magnitude as a function of luminance, F(l)= ml + c, where F(l) is the peak response at luminance l, c is the intercept, and m is the slope. The slope m served as the measure of luminance sensitivity, with non-zero values indicating luminance-dependent responses and the magnitude of m reflecting response strength. Statistical comparisons between slopes as mentioned in the figure legends.

### Visual stimulation at different wavelengths

ΔF/F_0_ was computed using the mean fluorescence signal in response to 682 nm (red LED) stimulation as the baseline (F_0_). Maximum responses to individual LED stimuli were extracted for analysis. Statistical comparison between L2 and L3 responses across wavelengths as mentioned in the figure legends.

### RNA sequencing library preparation

Brains were dissociated into single cells with the Neural Tissue Dissociation Kit (P) (Miltenyi Biotec, Cat. No. 130-092-628). Enzyme mix 1 (1,900 µl Buffer Z and 50 µl Enzyme P) and enzyme mix 2 (20 µl Buffer Y and 10 µl Enzyme A) were prepared freshly and mixed by vortexing. PBS was removed from each brain sample, and both enzyme mixes were divided equally between all samples. Alternatively, brain tissue was digested with 1 mg/ml papain (Sigma-Aldrich, Cat. No. P4762) and 1 mg/ml collagenase I (Sigma-Aldrich, Cat. No. SCR103) at 30 °C for 40 min.

After digestion, the enzyme was removed and 200 µl PBS were added. Tissue was physically dissociated by repeated pipetting until no intact tissue or debris remained visible. The suspension was brought to 400 µl with PBS and passed through a cell strainer with pore size of 30 µm (pluriSelect, Cat. No. 43-50030-03). When the suspension did not pass by gravity, it was drawn through the mesh by gentle pipetting from the underside.

Cells were sorted on BD FACSAria III using gate settings on forward and side scatter to exclude debris and doublets, and the GFP fluorescence positive gate was defined using the non-fluorescent negative control. The GFP positive cells for each replicate were sorted directly into 1.5 ml tubes containing 300 µl TRIzol reagent (Thermo Fisher Scientific, Cat. No. 10296010). Sorted samples were mixed by vortexing and stored at −80 °C until RNA extraction.

Total RNA was extracted following the TRIzol manufacturer’s protocol, except the aqueous phase was purified with QIAGEN RNeasy MinElute kit (Cat. No. 742024) with DNase I digestion. RNA concentration and integrity were assessed with Qubit RNA HS Assay (Thermo Fisher Scientific, Cat. No. Q10120) and Agilent Bioanalyzer RNA 6000 Pico chip (Cat. No. 5067-1513), respectively.

Sequencing libraries were prepared from total RNA using the NEBNext Ultra II Directional RNA Library Prep Kit for Illumina (New England Biolabs, Cat. No. E7770) according to the manufacturer’s instructions.

### RNA sequencing data processing

Sequencing libraries were run in paired-end mode on an Illumina HiSeq 2500 or NextSeq 500 platform. Demultiplexing and FASTQ file conversion were performed for all libraries using bcl2fastq (v1.8.4/v2.19.1), and raw read quality was evaluated with FastQC (v0.11.9). All reads were aligned to the BDGP6 reference genome using Ensembl (release 84)^22^ annotations. Transcript abundances were quantified as transcripts per million (TPM) utilizing Kallisto (https://pachterlab.github.io/kallisto/). The resulting count matrices were imported into the R environment and annotated via the TxDb.Dmelanogaster.UCSC.dm6.ensGene database. To ensure robust results, low-coverage transcripts were filtered out prior to downstream analysis, retaining only genes that met a minimum count threshold in at least one replicate per experimental condition.

Normalization and differential expression calculations were performed using the DESeq2 R package. To evaluate sample variance, identify potential batch effects, and detect outliers, Principal Component Analysis (PCA) was conducted on regularized log-transformed count data and visualized in 2D clustering plots. Significant differentially expressed genes (DEGs) in L3 vs L2 that we tested functionally were defined by an adjusted p-value of < 0.05 and a |log_2_ fold change| ≥ 3. The DEGS were graphically represented using volcano plots, mapping the log_2_ fold changes against their corresponding -log_10_ adjusted p-values. Finally, the Bioconductor org.Dm.eg.db database was used to map FlyBase gene identifiers ^23^ to their respective gene symbols, names, and Entrez IDs. To functionally annotate the DEGs, Gene Ontology (GO) enrichment analysis was performed utilizing the clusterProfiler package, which classified the data across three primary ontologies: Biological Process (BP), Molecular Function (MF), and Cellular Component (CC). The list of DEGs was subsequently refined by selecting for specific Gene Ontology classifications relevant to the study. Ion channels (GO:0005244, voltage-gated monoatomic ion channel activity) and neurotransmitter receptors (GO:003059, neurotransmitter receptor activity) (Fig. 7B) as well as ion channel regulators (GO:0099106, ion channel regulator activity) (Fig. 7C). Here, we considered all genes as differentially expressed genes (DEGs) in L3 vs L2 as well as regulated by dFezf if they satisfied adjusted p-value of < 0.05 and a |log2 fold change| ≥ 0.6 for both L2 vs. L3 expression and for L3 MARCM control vs. L3 MARCM *dFezf^1^* expression.

## Supplemental Figures and Figure legends

**Figure S1.**
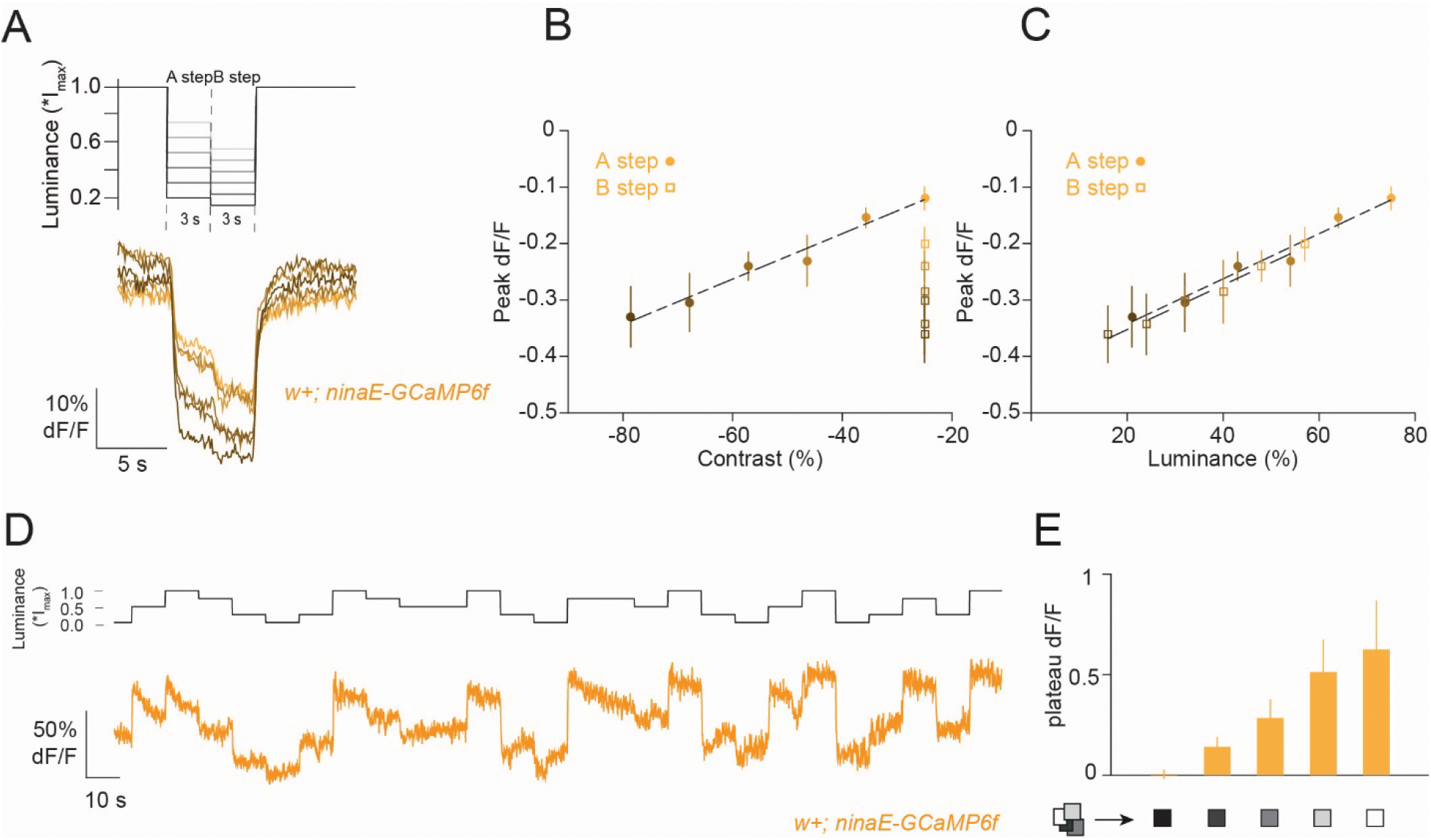
Photoreceptors R1-R6 encode both contrast and luminance. **(A)** Average calcium (GcaMP6f) signals from axon terminals of *ninaE-GCaMP6f* to the luminance contrast discrimination stimulus. **(B, C)** Peak responses (mean ± SEM) to the two OFF steps plotted as a function of contrast (**B**) and luminance (**C**). **(D)** Calcium signal trace of a single of *ninaE-GCaMP6f* to a stimulus of randomly varying luminance. The maximum screen luminance Imax was 1.87∗10^5^ photons∗s^−1^∗receptor^−1^. **(E)** Plateau responses (last 2 s of the epoch) of all steps for a given luminance. Sample size: n=7 (61) for (**A-C**) and 10 (89) for (**E**). Sample sizes are given as number of flies (number of cells).

**Figure S2.**
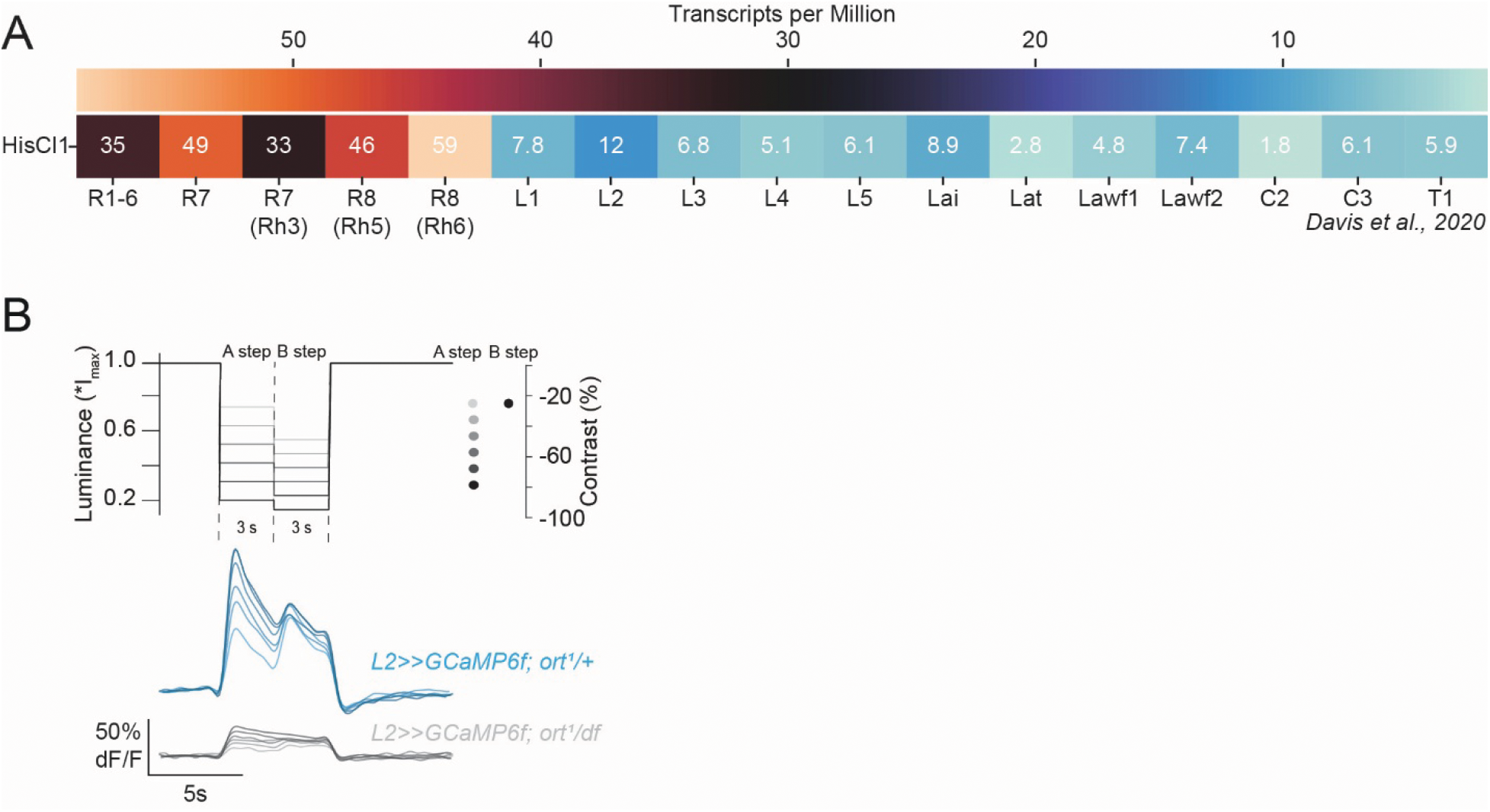
L2 neurons show a residual response in the absence of *ort*. **(A)** Heatmap of transcriptomic data of HisCl1 showing transcripts per million (TPM) for photoreceptors and lamina neurons obtained from the cell type-specific TAPIN-seq (GEO accession number GSE116969, Davis et al., 2020). **(B)** Average calcium (GCaMP6f) signals from axon terminals of L2 in *ort* mutant (, *L2>>GcaMP6f; ort^1^/df*) and *ort* heterozygous control ( *L2>>GcaMP6f; ort^1^/+*) to the luminance contrast discrimination stimulus. Sample size*, L2>>GCaMP6f; ort¹/+*: n=6 (56), *L2>>GCaMP6f; ort¹/df*: n=8 (66). Sample sizes are given as number of flies (number of cells).

**Figure S3.**
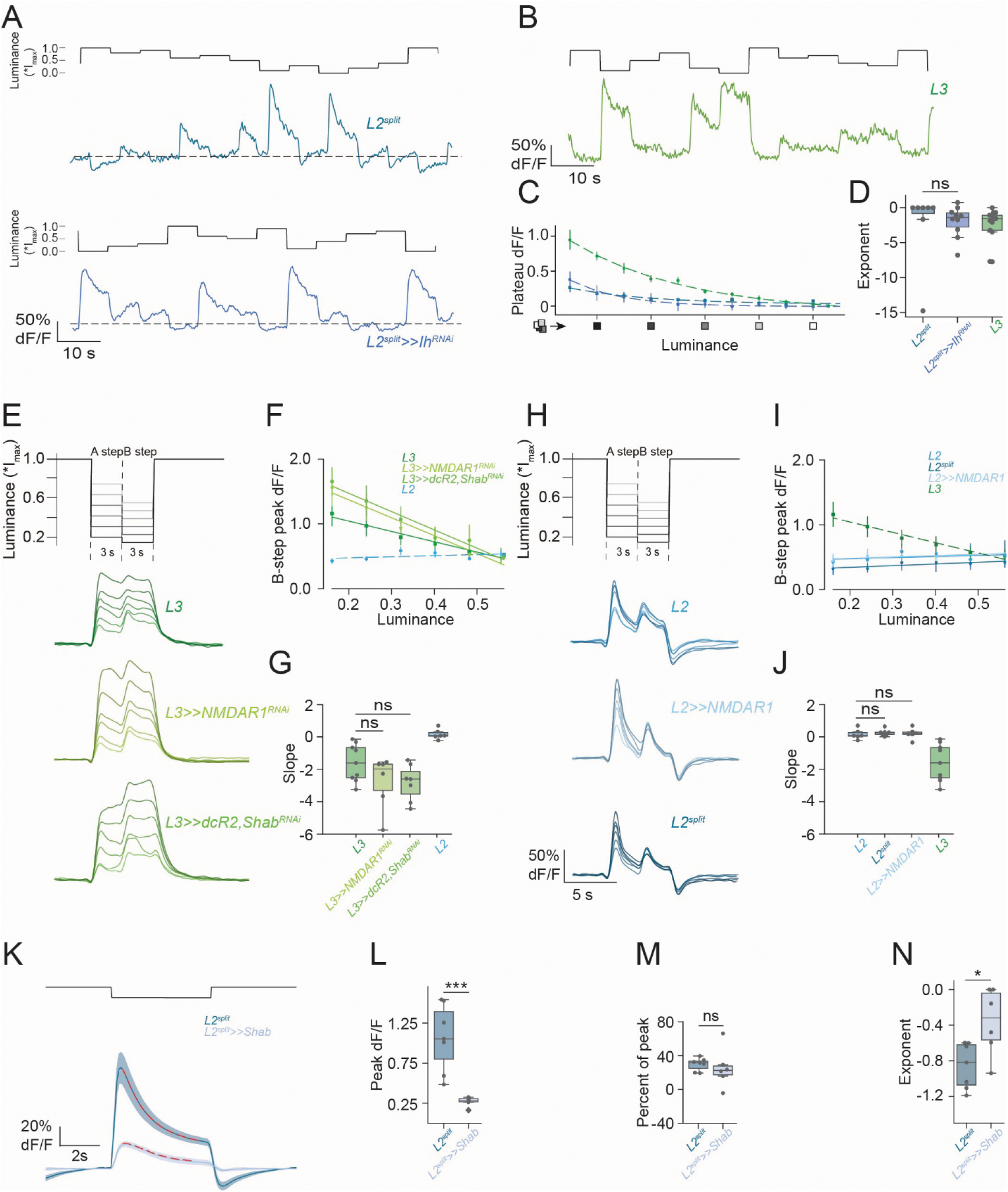
Manipulation of individual L2 enriched ion channels and receptors does not change L2 contrast and L3 luminance sensitivity. **(A-B)** Calcium response of a single of *L2^split^* control, *L2^split^>>Ih^RNAi^* and *L3* axon terminal to a stimulus of randomly varying luminance. The maximum screen luminance Imax was 1.<u>9</u>3∗10^5^ photons∗s^−1^∗receptor^−1^. **(C)** Plateau responses (last 500 ms of the epoch) of all steps for a given luminance. **(D)** Quantification of exponents from exponential fits to the plateau responses as a function of luminance. **(E)** Schematic of the luminance-contrast discrimination stimulus. Average calcium signals (GCaMP6f) from axon terminals of *L3* (green)*, L3>>NMDAR1^RNAi^, L3>>dcR2,Shab^RNAi^* to the luminance contrast discrimination stimulus. **(F)** Quantified mean peak ± SEM responses to B steps plotted as a function of luminance. **(G)** Slopes from linear fit to the responses in (F). **(H-J)** Same as (**E-G**) but for *L2* and *L2^split^ control, L2>>NMDAR1*. **(K)** Average calcium signals (GCaMP6f) for *L2^spli^control* neurons and *L2^split^>>Shab* to 5 s full field flashes. **(L)** Quantification for peak response, **(M)** Quantification for percentage of peak response retained during the plateau phase. **(N)** Quantification of exponents derived from exponential fits to OFF-step responses (red dotted line in (**K**), 1-4 s post-OFF step), ns, not significant, ***p<0.05, tested with two-tailed Student t test (**L**), *p<0.01, Mann-Whitney test, two-tailed (**N**). Sample size, *L2^split^*: n=7 (86)*, L2^split^>>Ih^RNAi^*: n=10 (98), *L3: n*=13 (114) for (**A-D**), *L2*: n=6 (66)*, L2^split^*: n=7 (85)*, L2>>NMDAR1*: n=6 (47), *L3: n*=9 (56)*, L3>>NMDAR1^RNAi^*: n=8 (164)*, L3>>dcR2,Shab^RNAi^*: n=7 (67) for (**E-J**), *L2^split^*: n=7 (88)*, L2^split^>>Shab*: n=6 (49) for (**K-N**).

**Figure S4.**
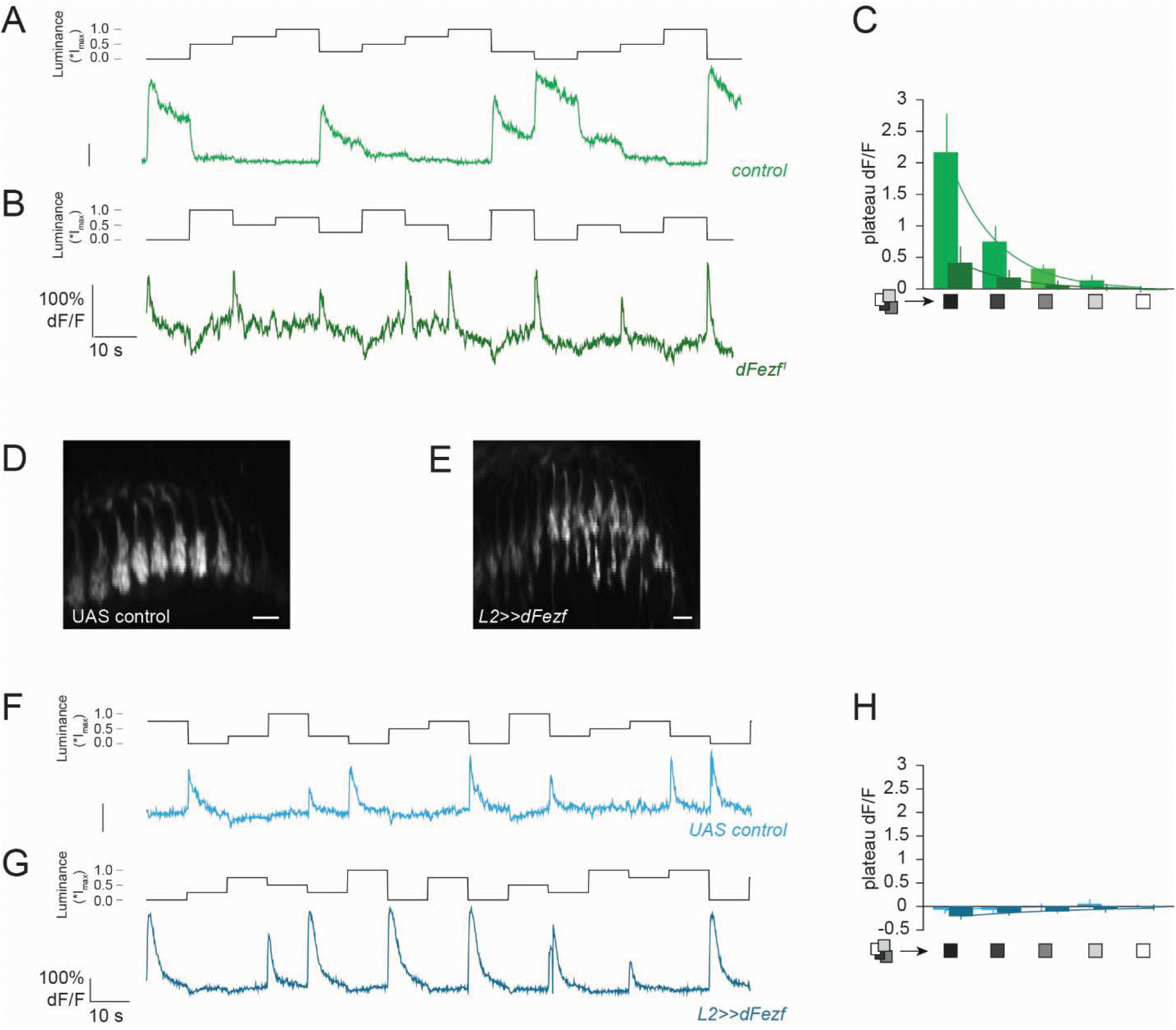
DFezf is not sufficient to transform L2 into luminance sensitive neuron. **(A,B)** Calcium response of a single of control L3 (**A**) and *dFezf-*mutant L3 (**B**) MARCM clone axon terminal to a stimulus of randomly varying luminance. The maximum screen luminance Imax was 1.87∗10^5^ photons∗s^−1^∗receptor^−1^. **(C)** Plateau responses (last 2 s of the epoch) of all step for a given luminance. **(D, E)** Representative in vivo calcium image of *UAS control L2* (**D**) or *L2>>dFezf* (**E**), dominantly marked by the expression of GCaMP6f. Scale bars are 10 μm. **(F, G)** Calcium response of a single of (F) UAS control L2 or (G) *L2>>dFezf* axon terminal to a stimulus of randomly varying luminance. The maximum screen luminance Imax was 1.87∗10^5^ photons∗s^−1^∗receptor^−1^. **(H)** Plateau responses (last 2 s of the epoch) of all steps for a given luminance. Sample size, Control L3: n=8 (21), *dFezf* mutant L3: n=12 (30) for (**C**), UAS Control L2: n=4 (60), L2>>*dFezf*: n=3 (40) for (**H**) Sample sizes are given as number of flies (number of cells).

## Bibliography

1. Baden, T., Berens, P., Franke, K., Román Rosón, M., Bethge, M., and Euler, T. (2016). The functional diversity of retinal ganglion cells in the mouse. Nature 529, 345–350. 10.1038/nature16468.

2. Franke, K., Berens, P., Schubert, T., Bethge, M., Euler, T., and Baden, T. (2017). Inhibition decorrelates visual feature representations in the inner retina. Nature 542, 439–444. 10.1038/nature21394.

3. Baden, T., Berens, P., Bethge, M., and Euler, T. (2013). Spikes in mammalian bipolar cells support temporal layering of the inner retina. Curr Biol 23, 48–52. 10.1016/j.cub.2012.11.006.

4. Wernet, M.F., and Silies, M. (2026). Imprecision in Vision: Lessons From Neural Circuits in the Fly. BioEssays 48, e70122. 10.1002/bies.70122.

5. Currier, T.A., Pang, M.M., and Clandinin, T.R. (2023). Visual processing in the fly, from photoreceptors to behavior. Genetics 224, iyad064. 10.1093/genetics/iyad064.

6. Ramos-Traslosheros, G., Henning, M., and Silies, M. (2018). Motion detection: Cells, circuits and algorithms. Neuroforum 24, A61–A72. 10.1515/nf-2017-A028.

7. Borst, A., and Groschner, L.N. (2023). How Flies See Motion. Annual Review of Neuroscience 46, 17–37. 10.1146/annurev-neuro-080422-111929.

8. Ramirez, L., Gjorgjieva, J., and Silies, M. (2026). Functional temporal cell type diversity in Drosophila reflects natural motion statistics. Preprint at bioRxiv, 10.64898/2026.04.30.721922 https://doi.org/10.64898/2026.04.30.721922.

9. Oesch, N.W., and Diamond, J.S. (2011). Ribbon synapses compute temporal contrast and encode luminance in retinal rod bipolar cells. Nature Neuroscience 14, 1555–1561. 10.1038/nn.2945.

10. Ketkar, M.D., Sporar, K., Gür, B., Ramos-Traslosheros, G., Seifert, M., and Silies, M. (2020). Luminance Information Is Required for the Accurate Estimation of Contrast in Rapidly Changing Visual Contexts. Current Biology 30, 657–669.e4. 10.1016/j.cub.2019.12.038.

11. Silies, M., Gohl, D.M., Fisher, Y.E., Freifeld, L., Clark, D.A., and Clandinin, T.R. (2013). Modular Use of Peripheral Input Channels Tunes Motion-Detecting Circuitry. Neuron. 10.1016/j.neuron.2013.04.029.

12. Fisher, Y.E., Leong, J.C.S., Sporar, K., Ketkar, M.D., Gohl, D.M., Clandinin, T.R., and Silies, M. (2015). A Class of Visual Neurons with Wide-Field Properties Is Required for Local Motion Detection. Current Biology 25, 3178–3189. 10.1016/j.cub.2015.11.018.

13. Euler, T., Haverkamp, S., Schubert, T., and Baden, T. (2014). Retinal bipolar cells: elementary building blocks of vision. Nat Rev Neurosci 15, 507–519. 10.1038/nrn3783.

14. Meinertzhagen, I.A., and O’Neil, S.D. (1991). Synaptic organization of columnar elements in the lamina of the wild type in Drosophila melanogaster. Journal of Comparative Neurology 305, 232– 263. 10.1002/cne.903050206.

15. Clark, D.A., Bursztyn, L., Horowitz, M.A., Schnitzer, M.J., and Clandinin, T.R. (2011). Defining the Computational Structure of the Motion Detector in Drosophila. Neuron 70, 1165–1177. 10.1016/j.neuron.2011.05.023.

16. Ketkar, M.D., Gür, B., Molina-Obando, S., Ioannidou, M., Martelli, C., and Silies, M. (2022). First-order visual interneurons distribute distinct contrast and luminance information across ON and OFF pathways to achieve stable behavior. eLife 11, e74937. 10.7554/eLife.74937.

17. Ketkar, M.D., Shao, S., Gjorgjieva, J., and Silies, M. (2023). Multifaceted luminance gain control beyond photoreceptors in Drosophila. Current Biology 33, 2632–2645.e6. 10.1016/j.cub.2023.05.024.

18. Gür, B., Ramirez, L., Cornean, J., Thurn, F., Molina-Obando, S., Ramos-Traslosheros, G., and Silies, M. (2024). Neural pathways and computations that achieve stable contrast processing tuned to natural scenes. Nat Commun 15, 8580. 10.1038/s41467-024-52724-5.

19. Puller, C., Ivanova, E., Euler, T., Haverkamp, S., and Schubert, T. (2013). OFF bipolar cells express distinct types of dendritic glutamate receptors in the mouse retina. Neuroscience 243, 136–148. 10.1016/j.neuroscience.2013.03.054.

20. DeVries, S.H. (2000). Bipolar Cells Use Kainate and AMPA Receptors to Filter Visual Information into Separate Channels. Neuron 28, 847–856. 10.1016/S0896-6273(00)00158-6.

21. DeVries, S.H., and Schwartz, E.A. (1999). Kainate receptors mediate synaptic transmission between cones and ‘Off’ bipolar cells in a mammalian retina. Nature 397, 157–160. 10.1038/16462.

22. Schnaitmann, C., Haikala, V., Abraham, E., Oberhauser, V., Thestrup, T., Griesbeck, O., and Reiff, D.F. (2018). Color Processing in the Early Visual System of Drosophila. Cell 172, 318–318.e18. 10.1016/j.cell.2017.12.018.

23. Gengs, C., Leung, H.-T., Skingsley, D.R., Iovchev, M.I., Yin, Z., Semenov, E.P., Burg, M.G., Hardie, R.C., and Pak, W.L. (2002). The Target of Drosophila Photoreceptor Synaptic Transmission Is a Histamine-gated Chloride Channel Encoded byort (hclA). J. Biol. Chem. 277, 42113–42120. 10.1074/jbc.M207133200.

24. Pantazis, A., Segaran, A., Liu, C.-H., Nikolaev, A., Rister, J., Thum, A.S., Roeder, T., Semenov, E., Juusola, M., and Hardie, R.C. (2008). Distinct Roles for Two Histamine Receptors (hclA and hclB) at the Drosophila Photoreceptor Synapse. Journal of Neuroscience 28, 7250–7259. 10.1523/jneurosci.1654-08.2008.

25. Witte, I., Kreienkamp, H.J., Gewecke, M., and Roeder, T. (2002). Putative histamine-gated chloride channel subunits of the insect visual system and thoracic ganglion. Journal of Neurochemistry 83, 504–514. 10.1046/j.1471-4159.2002.01076.x.

26. Hardie, R.C., and Weckström, M. (1990). Three classes of potassium channels in large monopolar cells of the blowfly Calliphora vicina. Journal of Comparative Physiology A 167, 723–736. 10.1007/BF00189763.

27. Rusanen, J., and Weckström, M. (2016). Frequency-selective transmission of graded signals in large monopolar neurons of blowfly Calliphora vicina compound eye. Journal of Neurophysiology 115, 2052–2064. 10.1152/jn.00747.2015.

28. Gür, B., Sporar, K., Lopez-Behling, A., and Silies, M. (2020). Distinct expression of potassium channels regulates visual response properties of lamina neurons in Drosophila melanogaster. J Comp Physiol A 206, 273–287. 10.1007/s00359-019-01385-7.

29. Pang, M.M., Chen, F., Xie, M., Druckmann, S., Clandinin, T.R., and Yang, H.H. (2025). A recurrent neural circuit in Drosophila temporally sharpens visual inputs. Current Biology 35, 333–346.e6. 10.1016/j.cub.2024.11.064.

30. Goaillard, J.-M., and Marder, E. (2021). Ion Channel Degeneracy, Variability, and Covariation in Neuron and Circuit Resilience. Annu. Rev. Neurosci. 44, 335–357. 10.1146/annurev-neuro-092920-121538.

31. Albantakis, L., Bernard, C., Brenner, N., Marder, E., and Narayanan, R. (2024). The Brain’s Best Kept Secret Is Its Degenerate Structure. J. Neurosci. 44, e1339242024. 10.1523/JNEUROSCI.1339-24.2024.

32. Rathour, R.K., and Narayanan, R. (2019). Degeneracy in hippocampal physiology and plasticity. Hippocampus 29, 980–1022. 10.1002/hipo.23139.

33. Schneider, M., Bird, A.D., Gidon, A., Triesch, J., Jedlicka, P., and Cuntz, H. (2023). Biological complexity facilitates tuning of the neuronal parameter space. PLoS Comput Biol 19, e1011212. 10.1371/journal.pcbi.1011212.

34. Dahmen, D., Hutt, A., Indiveri, G., Kennedy, A., Lefebvre, J., Mazzucato, L., Motter, A.E., Narayanan, R., Payvand, M., Planert, H., et al. (2025). How heterogeneity shapes dynamics and computation in the brain. Neuron, S0896627325008967. 10.1016/j.neuron.2025.11.023.

35. O’Leary, T., Williams, A.H., Caplan, J.S., and Marder, E. (2013). Correlations in ion channel expression emerge from homeostatic tuning rules. Proc. Natl. Acad. Sci. U.S.A. 110. 10.1073/pnas.1309966110.

36. Harris, W.A., Stark, W.S., and Walker, J.A. (1976). Genetic dissection of the photoreceptor system in the compound eye of Drosophila melanogaster. The Journal of physiology 256, 415–439. 10.1113/jphysiol.1976.sp011331.

37. Belušič, G., Ilić, M., Meglič, A., and Pirih, P. (2016). A fast multispectral light synthesiser based on LEDs and a diffraction grating. Sci Rep 6, 32012. 10.1038/srep32012.

38. Salcedo, E., Huber, a, Henrich, S., Chadwell, L.V., Chou, W.H., Paulsen, R., and Britt, S.G. (1999). Blue- and green-absorbing visual pigments of Drosophila: ectopic expression and physiological characterization of the R8 photoreceptor cell-specific Rh5 and Rh6 rhodopsins. The Journal of neuroscience : the official journal of the Society for Neuroscience 19, 10716–10726.

39. Rister, J., Pauls, D., Schnell, B., Ting, C.-Y., Lee, C.-H., Sinakevitch, I., Morante, J., Strausfeld, N.J., Ito, K., and Heisenberg, M. (2007). Dissection of the Peripheral Motion Channel in the Visual System of Drosophila melanogaster. Neuron 56, 155–170. 10.1016/j.neuron.2007.09.014.

40. Davis, F.P., Nern, A., Picard, S., Reiser, M.B., Rubin, G.M., Eddy, S.R., and Henry, G.L. (2020). A genetic, genomic, and computational resource for exploring neural circuit function. Elife 9. 10.7554/eLife.50901.

41. Gisselmann, G., Pusch, H., Hovemann, B.T., and Hatt, H. (2002). Two cDNAs coding for histamine-gated ion channels in D. melanogaster. Nat Neurosci 5, 11–12. 10.1038/nn787.

42. Döring, F., Wischmeyer, E., Kühnlein, R.P., Jäckle, H., and Karschin, A. (2002). Inwardly Rectifying K+ (Kir) Channels in*Drosophila*. Journal of Biological Chemistry 277, 25554–25561. 10.1074/jbc.M202385200.

43. Dahal, G.R., Rawson, J., Gassaway, B., Kwok, B., Tong, Y., Ptáček, L.J., and Bates, E. (2012). An inwardly rectifying K+ channel is required for patterning. Development 139, 3653–3664. 10.1242/dev.078592.

44. Marder, E., and Goaillard, J.-M. (2006). Variability, compensation and homeostasis in neuron and network function. Nat Rev Neurosci 7, 563–574. 10.1038/nrn1949.

45. Tan, L., Zhang, K.X., Pecot, M.Y., Nagarkar-Jaiswal, S., Lee, P.T., Takemura, S.Y., McEwen, J.M., Nern, A., Xu, S., Tadros, W., et al. (2015). Ig Superfamily Ligand and Receptor Pairs Expressed in Synaptic Partners in Drosophila. Cell 163, 1756–1769. 10.1016/j.cell.2015.11.021.

46. Xu, C., Ramos, T.B., Rogers, E.M., Reiser, M.B., and Doe, C.Q. (2024). Homeodomain proteins hierarchically specify neuronal diversity and synaptic connectivity. eLife 12, RP90133. 10.7554/eLife.90133.

47. Xu, C., Sastry, R.G., Newstein, P., Delgado, R.N., Wagh, N., Silies, M., Cepko, C.L., and Doe, C.Q. (2025). Progenitor-Derived Regulatory Logic Generates Functionally Related Neuronal Subtypes. Preprint at bioRxiv, 10.1101/2024.09.27.615552 https://doi.org/10.1101/2024.09.27.615552.

48. Peng, J., Santiago, I.J., Ahn, C., Gur, B., Tsui, C.K., Su, Z., Xu, C., Karakhanyan, A., Silies, M., and Pecot, M.Y. (2018). Drosophila fezf coordinates laminar-specific connectivity through cell-intrinsic and cell-extrinsic mechanisms. eLife 7, e33962–e33962. 10.7554/eLife.33962.

49. Santiago, I.J., Zhang, D., Saras, A., Pontillo, N., Xu, C., Chen, X., Weirauch, M.T., Mistry, M., Ginty, D.D., Pecot, M.Y., et al. (2021). Drosophila Fezf functions as a transcriptional repressor to direct layer-specific synaptic connectivity in the fly visual system. PNAS 118. 10.1073/pnas.2025530118.

50. Sampson, M.M., Gschweng, K.M.M., Hardcastle, B.J., Bonanno, S.L., Sizemore, T.R., Arnold, R.C., Gao, F., Dacks, A.M., Frye, M.A., and Krantz, D.E. (2020). Serotonergic modulation of visual neurons in Drosophila melanogaster. PLOS Genetics 16, e1009003. 10.1371/journal.pgen.1009003.

51. Pagni, M., Haikala, V., Oberhauser, V., Meyer, P.B., Reiff, D.F., and Schnaitmann, C. (2021). Interaction of “chromatic” and “achromatic” circuits in Drosophila color opponent processing. Curr Biol 31, 1687–1698.e4. 10.1016/j.cub.2021.01.105.

52. Longden, K.D., Rogers, E.M., Nern, A., Dionne, H., and Reiser, M.B. (2023). Different spectral sensitivities of ON- and OFF-motion pathways enhance the detection of approaching color objects in Drosophila. Nat Commun 14, 7693. 10.1038/s41467-023-43566-8.

53. Wardill, T.J., List, O., Li, X., Dongre, S., McCulloch, M., Ting, C.Y., O’Kane, C.J., Tang, S., Lee, C.H., Hardie, R.C., et al. (2012). Multiple spectral inputs improve motion discrimination in the drosophila visual system. Science 336, 925–931. 10.1126/science.1215317.

54. Nern, A., Loesche, F., Takemura, S., Burnett, L.E., Dreher, M., Gruntman, E., Hoeller, J., Huang, G.B., Januszewski, M., Klapoetke, N.C., et al. (2025). Connectome-driven neural inventory of a complete visual system. Nature 641, 1225–1237. 10.1038/s41586-025-08746-0.

55. Matsliah, A., Yu, S., Kruk, K., Bland, D., Burke, A.T., Gager, J., Hebditch, J., Silverman, B., Willie, K.P., Willie, R., et al. (2024). Neuronal parts list and wiring diagram for a visual system. Nature 634, 166–180. 10.1038/s41586-024-07981-1.

56. Henning, M., Ketkar, M.D., Lüffe, T., Gohl, D.M., Clandinin, T.R., and Silies, M. (2026). Inhibitory columnar feedback neurons are involved in motion processing in Drosophila. eLife 14, RP108529. 10.7554/eLife.108529.

57. Seung, H.S. (2024). Interneuron diversity and normalization specificity in a visual system. Preprint, 10.1101/2024.04.03.587837 https://doi.org/10.1101/2024.04.03.587837.

58. Gonzalez-Suarez, A.D., Zavatone-Veth, J.A., Chen, J., Matulis, C.A., Badwan, B.A., and Clark, D.A. (2022). Excitatory and inhibitory neural dynamics jointly tune motion detection. Current Biology 32, 3659–3675.e8. 10.1016/j.cub.2022.06.075.

59. El-Danaf, R.N., Rajesh, R., and Desplan, C. (2023). Temporal regulation of neural diversity in Drosophila and vertebrates. Seminars in Cell & Developmental Biology 142, 13–22. 10.1016/j.semcdb.2022.05.011.

60. Peng, J., Santiago, I.J., Ahn, C., Gur, B., Tsui, C.K., Su, Z., Xu, C., Karakhanyan, A., Silies, M., and Pecot, M.Y. (2018). Drosophila Fezf coordinates laminar-specific connectivity through cell-intrinsic and cell-extrinsic mechanisms. eLife 7, e33962. 10.7554/eLife.33962.

61. O’Leary, T., Williams, A.H., Franci, A., and Marder, E. (2014). Cell Types, Network Homeostasis, and Pathological Compensation from a Biologically Plausible Ion Channel Expression Model. Neuron 82, 809–821. 10.1016/j.neuron.2014.04.002.

## Methods bibliography

1. Gohl, D.M., Silies, M.A., Gao, X.J., Bhalerao, S., Luongo, F.J., Lin, C.C., Potter, C.J., and Clandinin, T.R. (2011). A versatile in vivo system for directed dissection of gene expression patterns. Nature Methods 2011 8:3 *8*, 231–237. 10.1038/nmeth.1561.

2. Rister, J., Pauls, D., Schnell, B., Ting, C.Y., Lee, C.H., Sinakevitch, I., Morante, J., Strausfeld, N.J., Ito, K., and Heisenberg, M. (2007). Dissection of the Peripheral Motion Channel in the Visual System of Drosophila melanogaster. Neuron 56, 155–170. 10.1016/j.neuron.2007.09.014.

3. Timofeev, K., Joly, W., Hadjieconomou, D., and Salecker, I. (2012). Localized Netrins Act as Positional Cues to Control Layer-Specific Targeting of Photoreceptor Axons in Drosophila. Neuron 75, 80–93. 10.1016/J.NEURON.2012.04.037.

4. Silies, M., Gohl, D.M., Fisher, Y.E., Freifeld, L., Clark, D.A., and Clandinin, T.R. (2013). Modular Use of Peripheral Input Channels Tunes Motion-Detecting Circuitry. Neuron 79, 111–127. 10.1016/j.neuron.2013.04.029.

5. Nern, A., Zhu, Y., and Zipursky, S.L. (2008). Local N-Cadherin Interactions Mediate Distinct Steps in the Targeting of Lamina Neurons. Neuron 58, 34–41. 10.1016/J.NEURON.2008.03.022.

6. Schnaitmann, C., Haikala, V., Abraham, E., Oberhauser, V., Thestrup, T., Griesbeck, O., and Reiff, D.F. (2018). Color Processing in the Early Visual System of Drosophila. Cell 172, 318–330.e18. 10.1016/J.CELL.2017.12.018.

7. Chen, Y., Akin, O., Nern, A., Tsui, C.Y.K., Pecot, M.Y., and Zipursky, S.L. (2014). Cell-type-Specific Labeling of Synapses In Vivo through Synaptic Tagging with Recombination. Neuron 81, 280–293. 10.1016/J.NEURON.2013.12.021.

8. Peng, J., Santiago, I.J., Ahn, C., Gur, B., Tsui, C.K., Su, Z., Xu, C., Karakhanyan, A., Silies, M., and Pecot, M.Y. (2018). Drosophila fezf coordinates laminar-specific connectivity through cell-intrinsic and cell-extrinsic mechanisms. Elife 7. 10.7554/ELIFE.33962.

9. Weng, M., Golden, K.L., and Lee, C.Y. (2010). dFezf/Earmuff Maintains the Restricted Developmental Potential of Intermediate Neural Progenitors in Drosophila. Dev. Cell 18, 126–135. 10.1016/J.DEVCEL.2009.12.007.

10. Janssens, D.H., Komori, H., Grbac, D., Chen, K., Koe, C.T., Wang, H., and Lee, C.Y. (2014). Earmuff restricts progenitor cell potential by attenuating the competence to respond to self-renewal factors. Development 141, 1036–1046. 10.1242/DEV.106534.

11. Asteriti, S., Liu, C.H., and Hardie, R.C. (2017). Calcium signalling in Drosophila photoreceptors measured with GCaMP6f. Cell Calcium 65, 40–51. 10.1016/J.CECA.2017.02.006.

12. Bloomquist, B.T., Shortridge, R.D., Schneuwly, S., Perdew, M., Montell, C., Steller, H., Rubin, G., and Pak, W.L. (1988). Isolation of a putative phospholipase c gene of drosophila, norpA, and its role in phototransduction. Cell 54, 723–733. 10.1016/S0092-8674(88)80017-5.

13. Wernet, M.F., Velez, M.M., Clark, D.A., Baumann-Klausener, F., Brown, J.R., Klovstad, M., Labhart, T., and Clandinin, T.R. (2012). Genetic dissection reveals two separate retinal substrates for polarization vision in drosophila. Current Biology 22, 12–20. 10.1016/j.cub.2011.11.028.

14. Dahal, G.R., Rawson, J., Gassaway, B., Kwok, B., Tong, Y., Ptácek, L.J., and Bates, E. (2012). An inwardly rectifying K+ channel is required for patterning. Development (Cambridge) 139, 3653– 3664. 10.1242/dev.078592.

15. Hürkey, S., Niemeyer, N., Schleimer, J.H., Ryglewski, S., Schreiber, S., and Duch, C. (2023). Gap junctions desynchronize a neural circuit to stabilize insect flight. Nature 2023 618:7963 *618*, 118– 125. 10.1038/s41586-023-06099-0.

16. Lee, T., and Luo, L. (2001). Mosaic analysis with a repressible cell marker (MARCM) for Drosophila neural development. Trends Neurosci. 24, 251–254. 10.1016/S0166-2236(00)01791-4.

17. Peirce, J.W. (2009). Generating Stimuli for Neuroscience Using PsychoPy. Front. Neuroinform. 2. 10.3389/NEURO.11.010.2008.

18. Oesch, N.W., and Diamond, J.S. (2011). Ribbon synapses compute temporal contrast and encode luminance in retinal rod bipolar cells. Nat. Neurosci. 14, 1555–1561. 10.1038/nn.2945.

19. Ketkar, M.D., Sporar, K., Gür, B., Ramos-Traslosheros, G., Seifert, M., and Silies, M. (2020). Luminance Information Is Required for the Accurate Estimation of Contrast in Rapidly Changing Visual Contexts. Current Biology 30, 657–669.e4. 10.1016/j.cub.2019.12.038.

20. Belušič, G., Ilić, M., Meglič, A., and Pirih, P. (2016). A fast multispectral light synthesiser based on LEDs and a diffraction grating. Scientific Reports 2016 6:1 *6*, 32012-. 10.1038/srep32012.

21. Fisher, Y.E., Leong, J.C.S., Sporar, K., Ketkar, M.D., Gohl, D.M., Clandinin, T.R., and Silies, M. (2015). A Class of Visual Neurons with Wide-Field Properties Is Required for Local Motion Detection. Current Biology 25, 3178–3189. 10.1016/j.cub.2015.11.018.

22. Hoskins, R.A., Carlson, J.W., Wan, K.H., Park, S., Mendez, I., Galle, S.E., Booth, B.W., Pfeiffer, B.D., George, R.A., Svirskas, R., et al. (2015). The Release 6 reference sequence of the Drosophila melanogaster genome. Genome Res. 25, 445–458. 10.1101/GR.185579.114.

23. Öztürk-Çolak, A., Marygold, S.J., Antonazzo, G., Attrill, H., Goutte-Gattat, D., Jenkins, V.K., Matthews, B.B., Millburn, G., dos Santos, G., Tabone, C.J., et al. (2024). FlyBase: updates to the Drosophila genes and genomes database. Genetics 227. 10.1093/GENETICS/IYAD211.

